# NSUN2-dependent 5-methylcytosine Modification Regulates Influenza A virus Gene Expression and Genomic Packaging

**DOI:** 10.1101/2025.11.26.690652

**Authors:** Jing Hu, Shengqiang Jiang, Wanling You, Ruiwei Hao, Jianchao Li, ShuJuan Gong, Xiao xiao, Haiyan Zhao, Long Liu, Hongying Chen

**Author notes:** Correspondence: Hongying Chen; Long Liu, **Email:** (H.C.); (L.L.). These authors contributed equally to the work (Jing Hu, Shengqiang Jiang, Wanling You). **Author Contributions:** Conceptualization, H.C.; Methodology, H.C., J.H., S.J., L. L., and H.Z.; Formal Analysis, S.J., J.H., H.C., and L. L.; Investigation, J.H., S.J., W.Y., J.L., R.H., S.G, and X.X.; Writing-Original Draft, J.H., and H.C.; Writing–Review & Editing, all authors; Supervision, H.C., and L. L.; Funding Acquisition, H.C., and L. L. **Competing Interest Statement:** All authors declare that they have no conflict of interest.

## Abstract

Emerging evidence indicates that methyltransferase NSUN2 catalyzes 5-methylcytosine (m^5^C) modifications on various viral RNAs and plays important roles in viral biology. However, the regulatory roles of NSUN2 in influenza A virus (IAV) replication have not been elucidated. Here, we revealed that NSUN2 negatively regulated the viral RNA transcription and protein production by knocking out and over-expressing *NSUN2*. By m^5^C MeRIP-seq and RNA-BisSeq, NSUN2-dependent m^5^C sites were identified on both the plus and minus viral RNA strands. In *NSUN2-*KO cells, the m^5^C modification on vRNAs was reduced, resulting in the production of a large number of deficient interfering particles (DIPs) which had lower vRNA content, imbalanced genome fragments, abnormal morphology and reduced pathogenicity. Mutation of m^5^C sites at the 5’ and 3’ ends of PB2 vRNA interfered with the selective packaging of the 8 vRNA segments into virus particles, resulting in the formation of a variety of abnormally packaged virus particles. PB2-vRNA mutants also had reduced replication ability and pathogenicity in mice. Overall, these data demonstrate that the m^5^C residues catalyzed by NSUN2 are required on vRNAs for the proper assembly of infectious viral particles, suggesting the depletion of m^5^C modification as a potential strategy that can be utilized to attenuate IAV strains.

**Significance Statement:** Influenza A virus (IAV) infections pose a significant threat to global public health by causing substantial morbidity and mortality. The segmented nature of the IAV genome requires precise regulation of the genomic assembly to produce infectious progeny virus particles. In this study, we demonstrate that m^5^C modification is crucial for the proper assembly of infectious viral particles. The lowered level of m^5^C modifications on vRNAs leads to the generation of defective interfering particles with reduced replication capacity and pathogenicity. Our data shed new light on the selective packaging mechanism of IAV segmented genome and highlight a potential new strategy for attenuating IAV strains by targeting the m^5^C modification machinery.

## Introduction

Influenza virus infection is an important cause of severe morbidity and mortality in humans and animals. Influenza A viruses (IAVs) in particular have caused four pandemics in human population in 1918, 1957, 1968 and 2009 (1, 2). Understanding the interplay between IAV and its hosts is critical for the development of effective approaches to prevent and treat the viral diseases. IAV belongs to *Orthomyxoviridae* family, and its genome consists of eight single-stranded negative-sense RNA segments (PB2, PB1, PA, HA, NP, NA, M, and NS) ranging from 0.9 to 2.3 kb in length (3). Each genomic viral RNA (vRNA) is bound by multiple copies of nucleoprotein (NP) and a heterotrimeric viral polymerase to form a viral ribonucleoprotein (vRNP) complex (4). The selectively packaged eight vRNPs in a typical IAV virion can be directly visualized by electron tomography (5–7) or multiple sequential fluorescence in situ hybridization (8). It has been revealed that the eight vRNPs are packaged in a “1+7” pattern, with a central vRNP surrounded by seven different vRNPs (7, 9, 10). In each of the eight vRNA segments, packaging signals have been identified in both the 5’ and 3’ non-coding regions (NCRs) and the adjacent terminal coding regions (11–16). Accumulating evidence suggests that packaging signals, vRNA-NP interactions, and vRNA-vRNA intersegment interactions mediated by the packaging signals and other regions play important roles in the selective genome packaging to form the specific “1+7” complex. However, it remains elusive how the eight unique vRNPs are selectively assembled into a progeny virion during IAV replication.

RNA modifications are widely distributed in different RNA molecules. To date, more than 100 types of chemical modifications have been found in all kinds of RNAs (17). Among them, 5-methylcytosine (m^5^C), in which RNA methylation occurs on the fifth carbon of cytosine, is one of the prevalent RNA modifications (18). The m^5^C modification regulates a wide range of cellular processes such as nuclear RNA export (19), mRNA stability (20, 21), mRNA translation (22) and innate immune response to virus infection (23, 24). 5-methylcytosine modification is catalyzed by m^5^C methyltransferases (writers). In eukaryotic cells, NOL1/NOP2/SUN domain (NSUN) family proteins and DNA methyltransferase (DNMT) h omolog DNMT2 have been identified as the m^5^C writers (25, 26). Among these writers, NSUN2 has been well characterized as a major m^5^C methyltransferase responsible for mRNA methylation (24, 27, 28). The m^5^C modification on RNA can be recognized by m^5^C-binding proteins (readers) including Aly/REF export factor (ALYREF) (19),Y-box-binding protein 1 (YBX1) (21) and YTH domain-containing family 2 (YTHDF2) (29). To remove the modification from RNA strand, m^5^C residues can be converted to 5-hydroxymethyl, 5-formyl, and 5-carboxylcytosine (hm^5^C, f^5^C and ca^5^C) by TET (ten–eleven translocation) family proteins (30) or Alpha-ketoglutarate-dependent dioxygenase ABH1 (ALKBH1) (31).

In virological researches, the discoveries of m^5^C methylation on Sindbis virus (32, 33) and adenovirus (34) mRNAs date back to 1970s. However, due to technological limitations at that time, the biological functions of the modification were not revealed. In recent years, m^5^C modification on viral RNAs of some DNA viruses (Hepatitis B virus (35–37) and Epstein-Barr virus (38)), retroviruses (Human Immunodeficiency Virus (27) and Murine Leukemia Virus (39, 40)) and positive-sense RNA viruses including Zika virus, Dengue virus, hepatitis C virus, poliovirus, porcine reproductive and respiratory syndrome virus, enterovirus 71 and coronavirus have been mapped (41–46). NSUN2-mediated m^5^C modification of HBV mRNA is involved in regulating reverse transcription, nuclear export, mRNA stability and particle production (35–37). The genomes of retroviruses (HIV-1 and MLV) are found to be highly enriched with m^5^C methylation. Retroviruses recruit host’s methyltransferase NSUN2 to add m^5^C to the viral transcripts, which is beneficial for the viral protein production and virus replication (27, 39, 40). NSUN2-dependent m^5^C methylated cytosines are distributed across the positive-sense genomic RNA of SARS-CoV-2. Disruption of the m^5^C machinery enhances viral replication and the virulence (46). In addition, it has also been reported that the expression of NSUN2, which negatively regulates type I interferon responses during various virus infections, is inhibited upon various virus infection to boost antiviral responses for the elimination of the viruses (23, 24).

In this study, we profiled the widespread distribution of m^5^C methylation sites in plus/minus RNA strands of IAV isolate A/WSN/33 (H1N1), by comparing the m^5^C Methylated RNA Immunoprecipitation Sequencing (MeRIP-seq) and RNA Bisulfide Sequencing (RNA-BisSeq) results of IAV RNAs produced in *NSUN2*-KO cells with those from the wild-type cells. Low-m^5^C IAVs were produced in *NSUN2*-KO cells or generated by site-directed mutagenesis, and their replication ability in cultured cells and pathogenicity in mice were examined. Our data suggest that IAV has evolved to acquire m^5^C modification to facilitate the selective packaging of its fragmented genome.

## Results

### NSUN2 negatively regulates IAV gene expression by modulating transcription

NSUN2 is involved in regulating the replication of various viruses, such as HIV, HBV, EV71, and SARS-CoV-2. However, whether it also modulates the replication cycle of IAV remains unknown. Here, we examined the expression levels of NSUN2 during IAV infection. NSUN2 mRNA level decreased at the early stage of infection, but it gradually increased with the extension of infection time and reached about 5 fold higher than the level in uninfected cells (*SI Appendix*, Fig. S1*A*), suggesting that NSUN2 might play regulatory roles in IAV infection. In order to study the function of NSUN2, we generated an *NSUN2* knockout (*NSUN2-*KO) A549 cell line using the CRISPR-Cas9 system. The gene knockout was confirmed by DNA sequencing (47). The loss of NSUN2 protein, which was verified by Western blot (*SI Appendix*, Fig. S1*B*), did not change the growth curve (*SI Appendix*, Fig. S1*C*) and morphology of A549 cells (*SI Appendix*, Fig. S1*D*).

To explore whether NSUN2 is involved in regulating IAV gene expression, the protein levels of HA and NP in A549 cells and *NSUN2*-KO cells at different time points of infection and infected at different MOI of 0.1 and 0.5 were detected (Fig. 1*A*). It is evident that the production of viral proteins was increased in *NSUN2-*KO cells, no matter the cells were in early or late infection, were infected with IAV at MOI of 0.1 or 0.5. By RT-qPCR, it revealed that most IAV mRNAs had higher levels in the *NSUN2*-KO cells than in A549 cells, except for PA and NA which did not show significant differences (Fig. 1*B*). In addition, we performed time-course measurement of NP RNAs (vRNA, mRNA and cRNA) during IAV infection, and the results showed that NSUN2 depletion elevated not only the mRNA synthesis but also the vRNA and cRNA levels (Fig. 1*C*).

**Figure 1.**
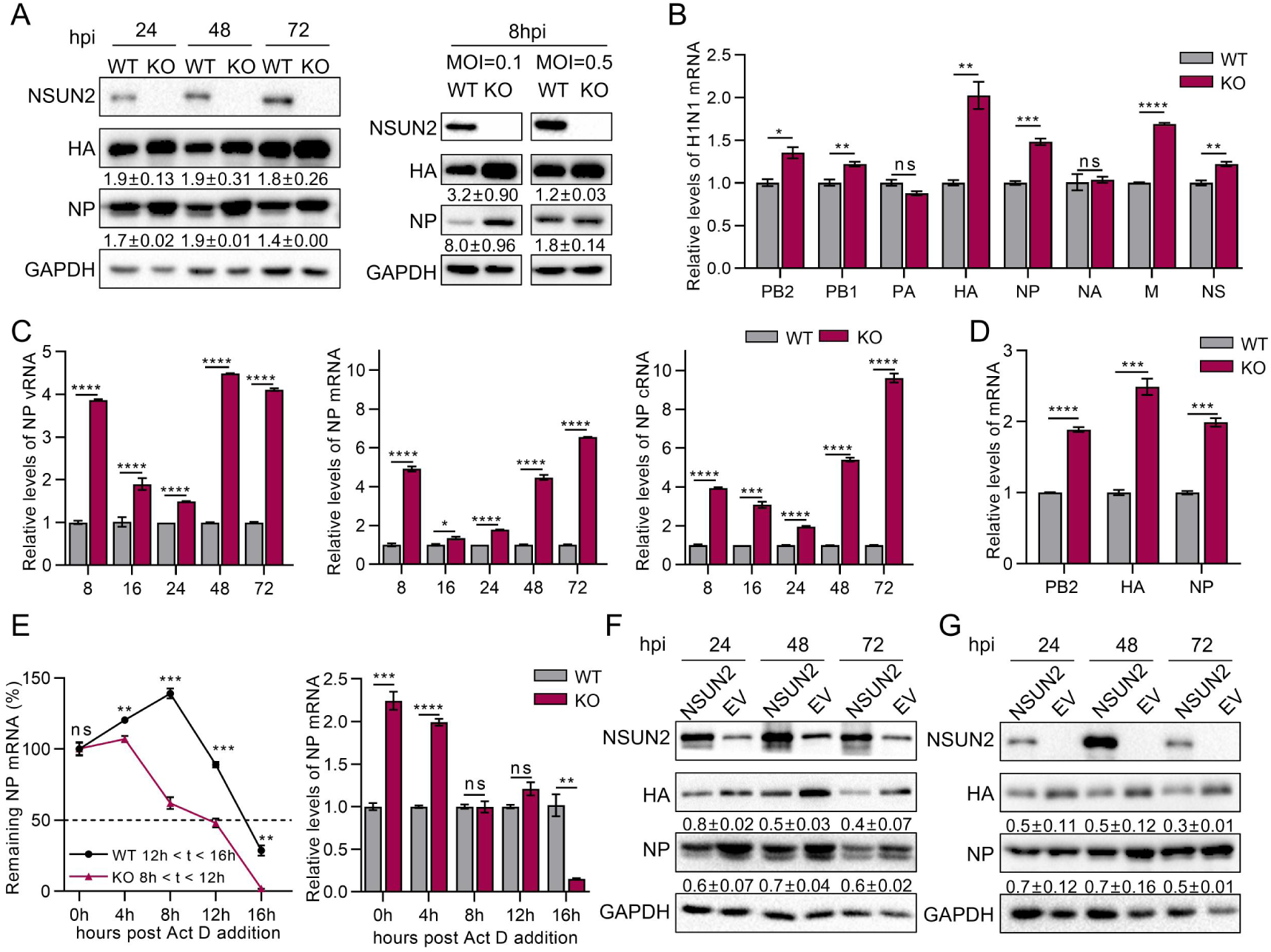
NSUN2 negatively regulates IAV gene expression. (A) Western blot detection of HA and NP protein levels in A549 (WT) and *NSUN2*-KO (KO) cells. The cells were infected with IAV-WSN at an MOI of 0.1 and harvested at 24, 48 and 72 hpi, or the cells were infected at an MOI of 0.1 or 0.5 and harvested at 8 hpi. (B) Quantification of IAV mRNA levels in A549 and *NSUN2-*KO cells by RT-qPCR. Cells were infected with IAV-WSN at an MOI of 0.1 and harvested at 36 hpi. (C) Quantification of IAV NP vRNA, mRNA and cRNA levels in A549 and *NSUN2*-KO cells by RT-qPCR. The cells were infected with IAV-WSN at an MOI of 0.1, and harvested at the indicated time points after infection. (D) *NSUN2* deletion enhances IAV primary transcription. Cells were infected with IAV at an MOI of 5 in the presence of 100 μg/mL cycloheximide (CHX). At 5 hpi, total RNAs were isolated for RT-qPCR analyses. (E) Examination of NP mRNA half-life. The cells were treated with actinomycin D (10 μg/mL) for 0, 4, 8, 12 and 16 h, and the mRNA levels were examined by RT-qPCR. The bar graph on the right shows the relative levels of NP mRNA at the indicated time points after actinomycin D treatment. (F) NSUN2 overexpression inhibits the production of HA and NP proteins in A549 cells. (G) Rescue expression of NSUN2 in *NSUN2*-KO cells suppresses IAV protein production. In (*F*) and (*G*), the cells were transfected with NSUN2 expressing plasmid (NSUN2) or empty vector (EV), then infected by IAV-WSN at 24 hpt, MOI=0.1. The relative protein expression levels between the WT and *NSUN2*-KO group (or the NSUN2 and EV group) were quantified by ImageJ scanning and normalized using GAPDH as the loading control. n = 3. Data are presented as mean ± SEM. ns, not significant, *, *P*<0.05, **, *P*<0.01, ***, *P*<0.001 and ****, *P*<0.0001.

In order to explore the mechanism by which NSUN2 regulates viral protein expression, we added cycloheximide (CHX), an inhibitor of eukaryotic protein synthesis, to block the synthesis of viral proteins in IAV infection. After the treatment with CHX, the synthesis of cRNA and the replication of vRNA are blocked due to the short of protein synthesis, but the primary viral mRNA transcription from the incoming vRNPs will still carry on (48, 49). Compared with the wild-type A549 cells, *NSUN2* deletion significantly promoted the primary transcription of IAV genes in the presence of CHX (Fig. 1*D*). Since NSUN2 has been reported to regulate the stability of RNAs, we examined NP mRNA half-life after the treatment of actinomycin D (Act D). The viral mRNA in *NSUN2*-KO cells obviously decayed faster than in the A549 cells (Fig. 1*E*, left panel). Despite having a shorter half-life, the viral mRNA had a higher level in *NSUN2*-KO cells than that in the wild-type cells till 8 hours after the addition of Act D (Fig. 1*E*, right panel). These results demonstrate that *NSUN2* deletion promoted viral gene expression by enhancing the transcriptional activity of vRNP but not the mRNA stability.

To verify the regulation role of NSUN2 in IAV gene expression, A549 cells and *NSUN2*-KO cells were transiently transfected with pcDNA3.1-NSUN2 to overexpress or rescue the methyltransferase, followed by the infection of WSN at an MOI of 0.1. Both overexpression of NSUN2 in A549 cells (Fig. 1*F*) and rescue expression of NSUN2 in *NSUN2-*KO cells (Fig. 1*G*) caused significant reduction of HA and NP protein levels, confirming the downregulation role of NSUN2 in IAV gene expression.

### NSUN2 positively regulates the production of IAV progeny virus

We next tested whether NSUN2 affected the production of progeny viruses. To assess the infectivity of viruses, we measured the hemagglutination (HA) titer and the plaque forming units (PFU) of IAVs released from the virus infected cells. A relative PFU/HA ratio was calculated to reflect the change of the infectious particles relative to the total virus particles produced (50). The progeny viruses released from *NSUN2-*KO cells had a lower viral titer and a higher HA titer, consequently a much lower PFU/HA ratio than the virus released from wild-type A549 cells (Fig. 2*A* and *SI Appendix*, Fig. S2*A*). When NSUN2 was overexpressed in A549 cells, it did not significantly change the production of infectious IAV but resulted in a lower HA titer and slightly increased PFU/HA ratio, compared with the empty vector group (Fig. 2*B* and *SI Appendix*, Fig. S2*B*). Rescue expression of NSUN2 in *NSUN2-*KO A549 cells restored the viral titer and PFU/HA ratio (Fig. 2*C* and *SI Appendix*, Fig. S2*C*). By plaque assay, we observed that compared to the viruses produced from the parental A549 cells, the viruses produced from the *NSUN2*-KO A549 cells formed obviously smaller plaques when they were used to infect MDCK cells (Fig. 2*D* and *SI Appendix*, Fig. S2*D*). Overexpression of NSUN2 did not apparently affect the plaque size (Fig. 2*E* and *SI Appendix*, Fig. S2*E*). However, when NSUN2 was rescue-expressed in the *NSUN2*-KO cells, the plaque size of viruses was restored to normal (Fig. 2*F* and *SI Appendix*, Fig. S2*F*). These data suggest that NSUN2 could facilitate the efficient production of infectious IAV particles.

**Figure 2.**
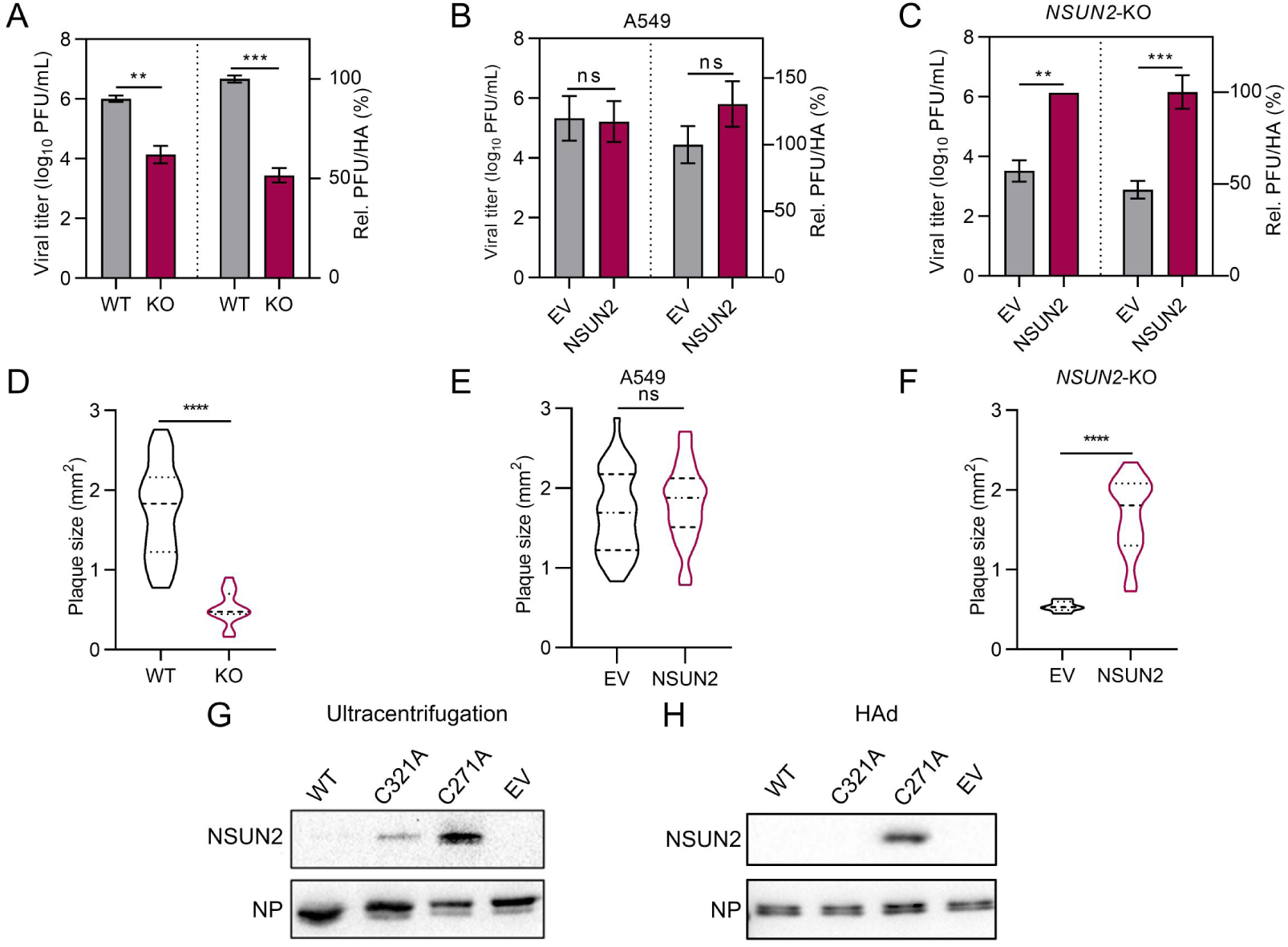
NSUN2 positively regulates the production of IAV progeny virus. (*A-C*) The influences of NSUN2 on IAV titer and PFU/HA ratio. The cell culture media from IAV infected A549 cells (WT) and *NSUN2-*KO A549 cells (KO) (*A*), A549 cells overexpressing NSUN2 (*B*) and *NSUN2-*KO cells rescue-expressing NSUN2 (*C*) were harvested at 72hpi. EV, cells transfected with empty vector. (*D-F*) Plaque sizes for the IAV stocks harvested in (*A*) to (*C*). The plaque images are shown in *SI Appendix*, Fig. S2*D-F*. Data are presented as mean ± SEM from n = 3. ****, *P*<0.0001 and ns, not significant by two tailed *t-test* between the indicated groups. (*G*) and (*H*) Western blot detection of NSUN2 packaged in IAV virus particles, which were purified by ultracentrifugation (*G*) or HAd (*H*) from the cell culture media of IAV-infected *NSUN2-*KO cells rescue-expressing NSUN2 (WT) or the indicated mutants. EV, cells transfected with empty vector as a negative control. IAV NP was examined as a control protein packed in virus particles. The cells were infected with IAV-WSN (*A*) or infected after the transfection of the indicated protein expression plasmids at 24 hpt (*B*-*H*). The cell culture supernatants were harvested for virus titration and purification at 72 hpi.

The methyltransferases in NSUN family contain two conserved cysteine, one of which (C321 in NSUN2) is responsible for forming a covalent bond to the pyrimidine base, while the other cysteine (C271 in NSUN2) is essential for the breakdown of the covalent linkage between the enzyme and the RNA molecule (51). Therefore, C321A mutation can result in an inactive enzyme, whereas C271A mutant can form stable crosslinks with RNA strands at the target cytosine sites (52). To investigate whether NSUN2 physically interacts with the vRNAs packaged in IAV particles, we transfect the plasmids expressing the NSUN2 mutants into the *NSUN2-*KO A549 cells, then infected the cells with IAV, and purified the virus particles released into the cell media by ultracentrifugation through a 20-60% sucrose gradient. The results showed that the NSUN2-C271A mutant was efficiently packaged into IAV (Fig. 2*G*). Meanwhile, a small amount of wild-type NSUN2 and NSUN2-C321A mutant were also detected in the virus particles, which may come from some cellular components retained in the purified virus sample. To avoid the contamination of cell debris and exextracellular vesicles in the influenza virus purification, the viruses were further purified by hemadsorption/elution assay (HAd) (53–55). In HAd, virus particles were captured by red blood cells through the receptor-binding activity of HA and released through the cleavage activity of NA (the protocol is illustrated in *SI Appendix*, Fig. S2*G*). The result showed that only NSUN2-C271A was detected in the purified virus particles (Fig. 2*H*). These results demonstrate the spontaneous crosslink of NSUN2-C271A to IAV particles and suggest a potential role of NSUN2 in adding m^5^C modification to IAV vRNAs.

### NSUN2 regulates the m^5^C modifications on IAV plus- and minus-strand RNAs

To investigate whether the IAV RNAs are modified by m^5^C methylation and NSUN2 regulates this modification, we conducted a transcriptome-wide RNA modification analysis of both the plus and minus RNA strands of IAV in the virus-infected A549 cells and *NSUN2*-KO cells. To estimate m^5^C modifications on IAV RNAs, IAV mRNA extracted from H1N1-infected cells at 36 hpi and vRNAs extracted from released virus particles at 60 hpi were analyzed by MeRIP-qPCR. The m^5^C modified RNA levels of most mRNA and vRNA fragments from *NSUN2-*KO cells were significantly lower than those from wild-type A549 cells, except that PA had a higher level of modified mRNA after NSUN2 depletion, and no significant differences were detected on PA vRNA, NA mRNA and HA RNAs (*SI Appendix*, Fig. S3*A* and S3*B*). To map the m^5^C modification regions, the total RNA samples isolated from H1N1-infected cells at 36 hpi were subjected to m^5^C MeRIP-seq. In order to distinguish the IAV mRNA/cRNA (plus) and vRNA (minus) strands, we adopted strand-specific library construction to identify the orientation of the RNA segments. Cellular mRNAs for ENO1 (56), PFAS (57), ICAM1 (58), and HDGF (21), which have been reported to be methylated by NSUN2, were examined as positive controls in the analysis of MeRIP-seq data (*SI Appendix*, Fig. S3*C*). As shown in Fig. 3*A*, most of the IAV plus-strand (mRNA/cRNA) and minus-strand (vRNA) segments bear N5-methylcytosine modifications. There are abundant m^5^C peaks distributed on the plus-strand RNAs with no obvious distribution preference. On the vRNAs, some m^5^C peaks located in or close to the 3’ or 5’ UTRs, which are required for genome incorporation and virion assembly (59). In *NSUN2*-KO cells, most m^5^C peaks on both the IAV plus and minus strands are lost, compared with the RNAs extracted from A549 cells. However, a distinct new m^5^C peak appears on the PA plus-strand, and two peaks are retained on the PA minus-strand. These m^5^C residues on PA RNAs could be added by a different enzyme(s) in the *NSUN2*-KO cell.

**Figure 3.**
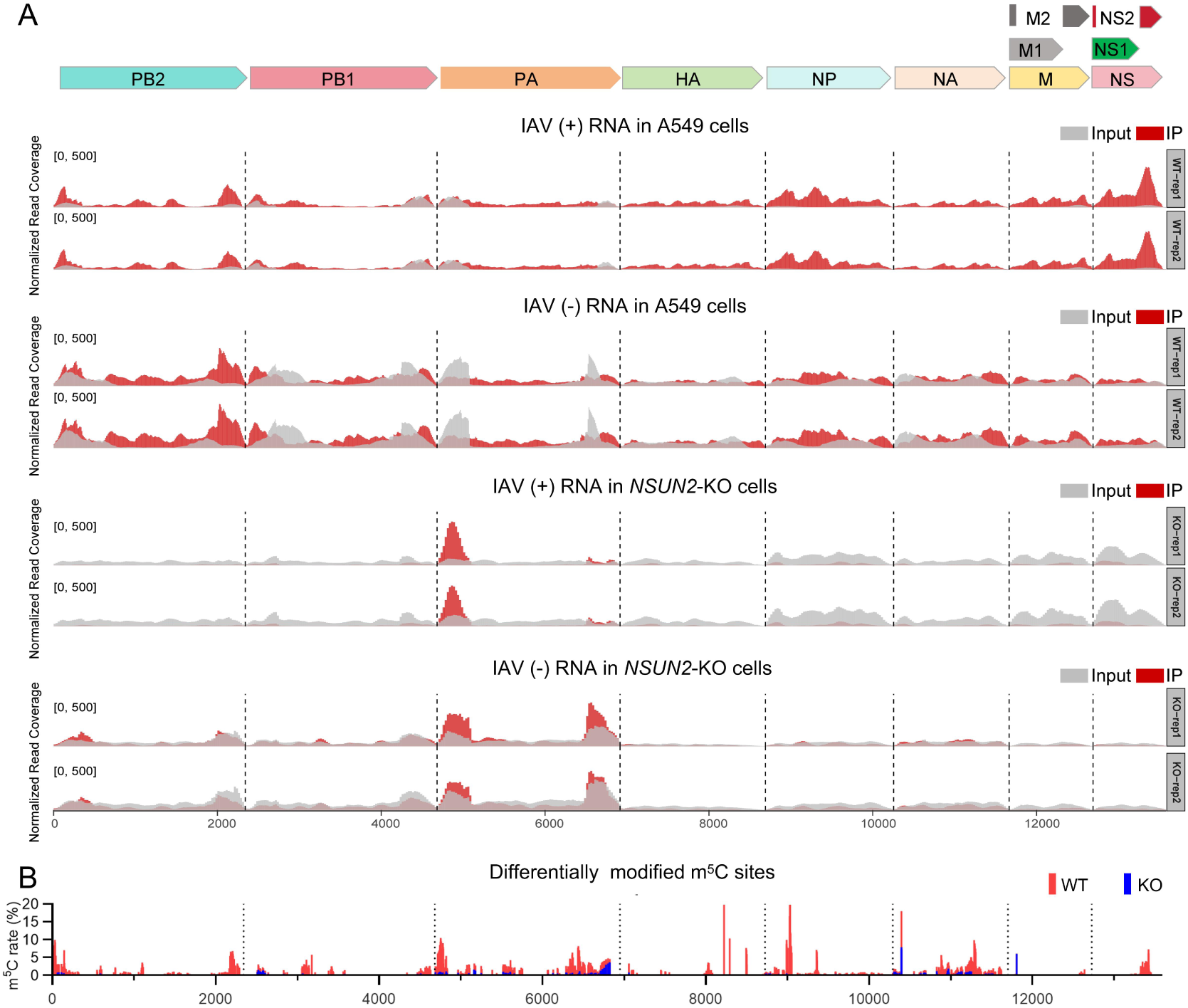
NSUN2 regulates the m^5^C modifications on IAV plus and minus RNA strands. (A) The m^5^C peaks on IAV plus and minus RNA strands identified by m^5^C MeRIP-seq. The concatenated map of the IAV-WSN genome is shown on the top for the alignment of detected methylation sites. The A549 cells or *NSUN2*-KO cells used for total RNA extraction were infected with IAV-WSN at an MOI of 0.1, and harvested at 36 hpi. Subsequently, the m^5^C peaks were detected by MeRIP-seq. Data for two independent experiments are shown. (B) Potential m^5^C modified sites on IAV vRNA genome identified by RNA-BisSeq. A549 cells and *NSUN2*-KO cells were infected with IAV-WSN at an MOI of 0.1. At 60 hpi, the virus particles released in the cell culture supernatant were used for RNA extraction and subjected to RNA-BisSeq. Differential sites identified between the vRNAs from WT and *NSUN2*-KO cells are presented here. The sequencing results for the two repeats of vRNAs respectively extracted from WT and *NSUN2*-KO A549 cells are shown in *SI Appendix*, Fig. S4.

In order to identify the specific m^5^C sites on the genomic vRNAs packaged in IAV virions, we concentrated IAV particles released in the supernatant of H1N1-infected wild-type A549 cells and *NSUN2*-KO cells at 60 hpi using PEG6000. The viral RNAs were isolated and subjected to RNA bisulfite sequencing (RNA-BisSeq, n = 2), which identified multiple sites having ≥ 1% C reads retained after bisulfite treatment on the vRNAs produced in A549 cells. Compared with the results detected in A549 cells, the potential modification sites on vRNAs produced in *NSUN2*-KO cells were significantly reduced (*SI Appendix*, Fig. S4). The m^5^C sites differentially identified of virion on the vRNAs from the WT group versus *NSUN2*-KO group were presented as the potential m^5^C modification sites catalyzed by NSUN2 (Fig. 3*B*). Overall, by MeRIP-seq and RNA-BisSeq, we found that both the plus- and minus- strands of IAV RNAs harbor multiple m^5^C modifcation sites. The absent of the majority of these m^5^C residues in the RNAs produced in *NSUN2*-KO cell suggest that these sites are mainly modified by NSUN2.

### The low-m^5^C IAV forms defective interfering particles with disordered vRNA packaging

Compared with the IAV in WT A549 cells, IAV replicated in *NSUN2*-KO cells has lower level of m^5^C modification (Fig. 3), and produces lower level of infectious virus particles (Fig. 2) but higher level of viral proteins (Fig. 1). To further characterize the low-m^5^C IAV produced in *NSUN2*-KO cells, we examined the abundance and ratio of vRNAs using RT-qPCR. In the cytoplasm of the virus infected cells, all eight vRNA segments were more abundantly produced in *NSUN*2-KO than in WT A549 cells, but the relative proportions of the vRNAs were similar between the two groups (Fig. 4*A*). In the virus particles released in the cell culture media, however, the relative abundance and proportion of PB2, PB1 and PA were significantly increased, while the other five segments were obviously decreased (Fig.4*B*). By Digital PCR, it was confirmed that knockout of *NSUN2* resulted in the release of much lower copies of genomic vRNAs (Fig. 4*C*). Therefore, we speculated that the loss of m^5^C modification on vRNAs in *NSUN2*-KO cells might have interfered with the interactions between RNA fragments or the interactions between RNA fragment and viral structural proteins during the virus assembly, and consequently resulted in some deficient virus particles. To verify this hypothesis, virus particles were observed by negative staining transmission electron microscopy (ns-TEM). The analysis detected 48.58% (n = 212) of empty virus-like particles and vRNP packaging-deficient particles containing obviously less than 8 genome fragments budding from *NSUN2*-KO cells. In comparison, the number of vRNP packaging-deficient particles only accounted for 23.43% (n = 303) of the virus particles budding from WT A549 cells (Fig. 4*D*).

**Figure 4.**
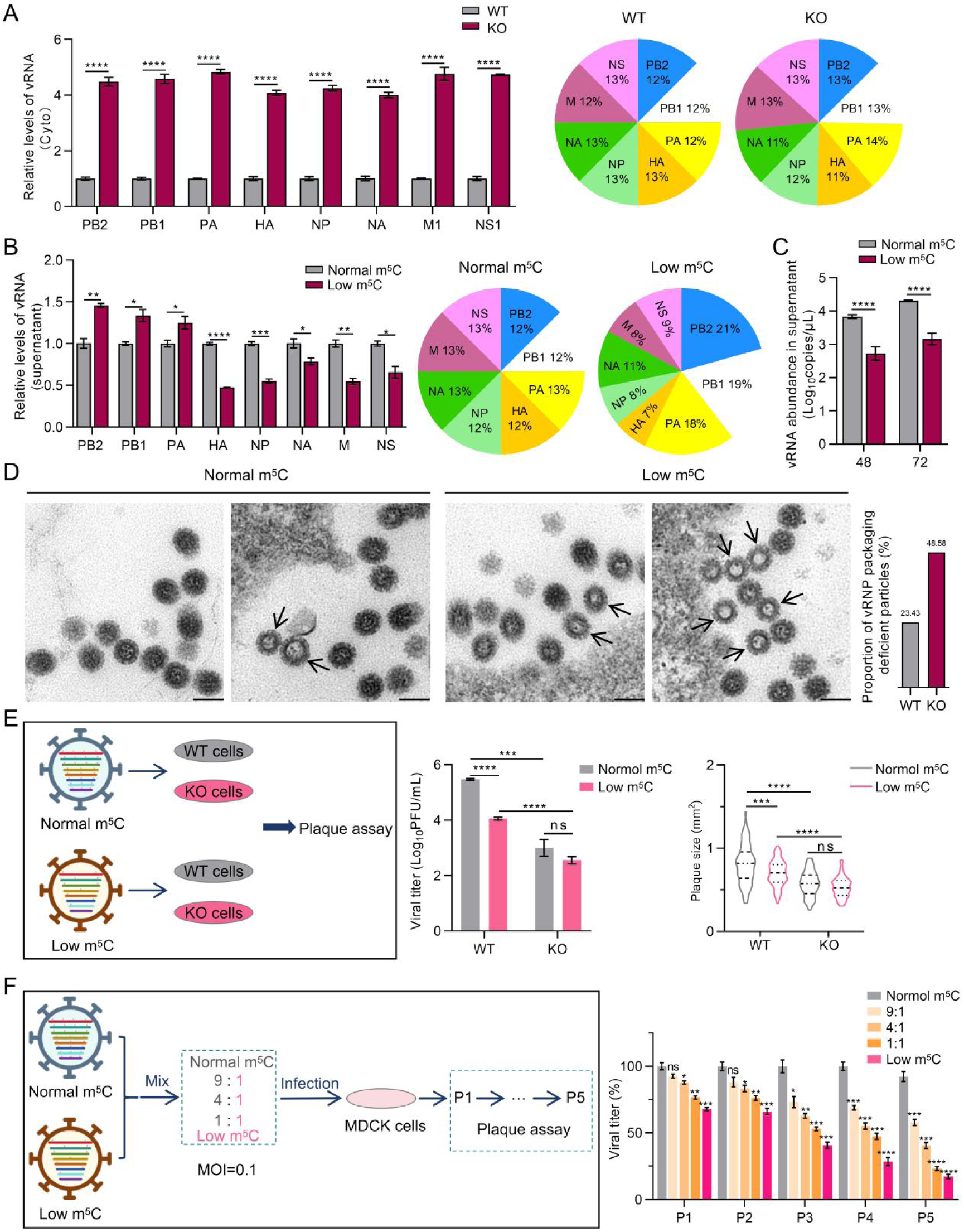
The low-m^5^C IAV forms vRNP packaging-deficient particles and acts as defective interfering particles. (A) Detection of IAV vRNA levels in the cytoplasm of *NSUN2-*KO A549 cells versus the WT A549 cells by RT-qPCR. Cells were infected with IAV-WSN at an MOI of 1, and the cytoplasmic RNAs were extracted at 6 hpi. The relative proportion of vRNAs are calculated based on the RNA levels and shown with the pie charts. (B) Detection of IAV vRNA levels in the cell culture supernatant of *NSUN2*-KO cells (Low m^5^C) versus the wild-type A549 cells (Normal m^5^C) by RT-qPCR. The cells were infected with IAV-WSN at an MOI of 0.1, and the supernatants were harvested at 60 hpi to extract RNA from viral particles. The relative proportion of genomic vRNAs in virus particles are calculated based on the RNA levels and shown with the pie charts. (C) Digital RT-PCR detection of total vRNA copies released into the cell culture media. WT and *NSUN2*-KO A549 cells were infected with IAV-WSN at an MOI of 0.1, and the vRNAs released at 48 and 72 hpi were examined. (D) Observation of the virus particles produced from A549 cells (Normal m^5^C) and *NSUN2*-KO cells (Low m^5^C) by negative staining transmission electron microscopy. Scale bar, 100 nm. Arrows indicate empty particles and vRNP packaging-deficient virus particles. The percentage of vRNP packaging-deficient virus particles is presented in a bar chart on the right. (E) Propagating and cross-passaging of IAV in WT and *NSUN2*-KO A549 cells. Flow chart for the experiment is shown in the left box. WT and KO cells were infected with IAV-WSN at an MOI of 0.1, producing virus stocks with normal m^5^C and low m^5^C. Then, WT and *NSUN2*-KO A549 cells were respectively infected with the normal-m^5^C and low-m^5^C viruses. The progeny viruses were harvested at 72 hpi for plaque assay on MDCK cells. Virus titers (the middle chart) and plaque sizes (the right chart) of the cross-passaged viruses formed on MDCK cell monolayers are quantified. (F) Co-infecting MDCK cells with normal and low m^5^C IAVs. Flow chart for the experiment is shown in the left box. The normal and low m^5^C viruses were mixed at ratios of 9:1, 4:1 and 1:1, or respectively used to infect MDCK cells, at a total MOI of 0.1 for each infection. The progeny viruses were continuously passaged on MDCK cells for five times. Virus titers of the indicated viruses through continuous passage are shown in a bar chart on the right. Data are presented as mean ± SEM (n=3). *, *P*<0.05, **, *P*<0.01, ***, *P*<0.001, ****, *P*<0.0001 and ns, not significant.

To explore whether the replication ability and plaque forming property of low-m^5^C IAV, which was harvested from *NSUN2*-KO cells with epigenetic defects of m^5^C modification, can be recovered by replication in WT cells expressing NSUN2, we crossly passaged the viruses harvested from the WT and *NSUN2*-KO A549 cells. As shown in Fig.4*E*, re-introducing the low-m^5^C virus into WT A549 cells neither restored the virus titer nor the plaque size.

With a disordered proportion of vRNPs in the virus particles, some low-m^5^C virus particles may act as defective interfering particles (DIPs) to interfere with the replication of standard viruses (STVs) (60–62). To explore this possibility, we co-infected MDCK cells with normal-m^5^C viruses and low-m^5^C viruses at a total MOI of 0.1. In comparison with the normal-m^5^C viruses, co-infection of MDCK cells resulted in progeny viruses at lower titers. The higher the proportion of low-m^5^C viruses in co-infection, the greater the decrease in virus titer. Continuous passage of the progeny viruses in MDCK cells did not restore the virus replication but expanded the differences in titer compared to the normal-m^5^C virus. After 5 passages, the titers of progeny viruses from the 1:1 co-infection group dropped to a level close to the continuously passaged low-m^5^C virus (Fig. 4*F*). These data indicate that the low-m^5^C virus is a type of DIP that can interfere the replication of normal-m^5^C virus.

### The low-m^5^C IAV has reduced replication ability and pathogenicity in mice

To examine the influence of m^5^C modification of vRNAs on the virulence of IAV *in vivo*, mice (n = 5) were infected with low-m^5^C IAV generated from *NSUN2-*KO cells in parallel with the wild-type IAV produced in A549 cells, at a dose of 1×10^5^ PFU. The body weight of infected mice decreased from the second day of infection and the survivals started to recover at 10 dpi. During this period, the survivals infected with low-m^5^C virus had reduced weight loss compared with the WT group (Fig.5*A*). In two weeks, all the mice infected by the WT virus died. In contrast, three of the five mice infected with low-m^5^C IAV survived (Fig.5*B*). At 5 dpi, the titers of WT virus in lungs were significantly higher than the low-m^5^C virus (Fig. 5*C*). Consistent with its lower replication ability, the low-m^5^C IAV infection caused milder pathological changes in lungs than the WT virus (Fig. 5*D*), demonstrating that the low-m^5^C IAV had reduced pathogenicity in mouse model.

**Figure 5.**
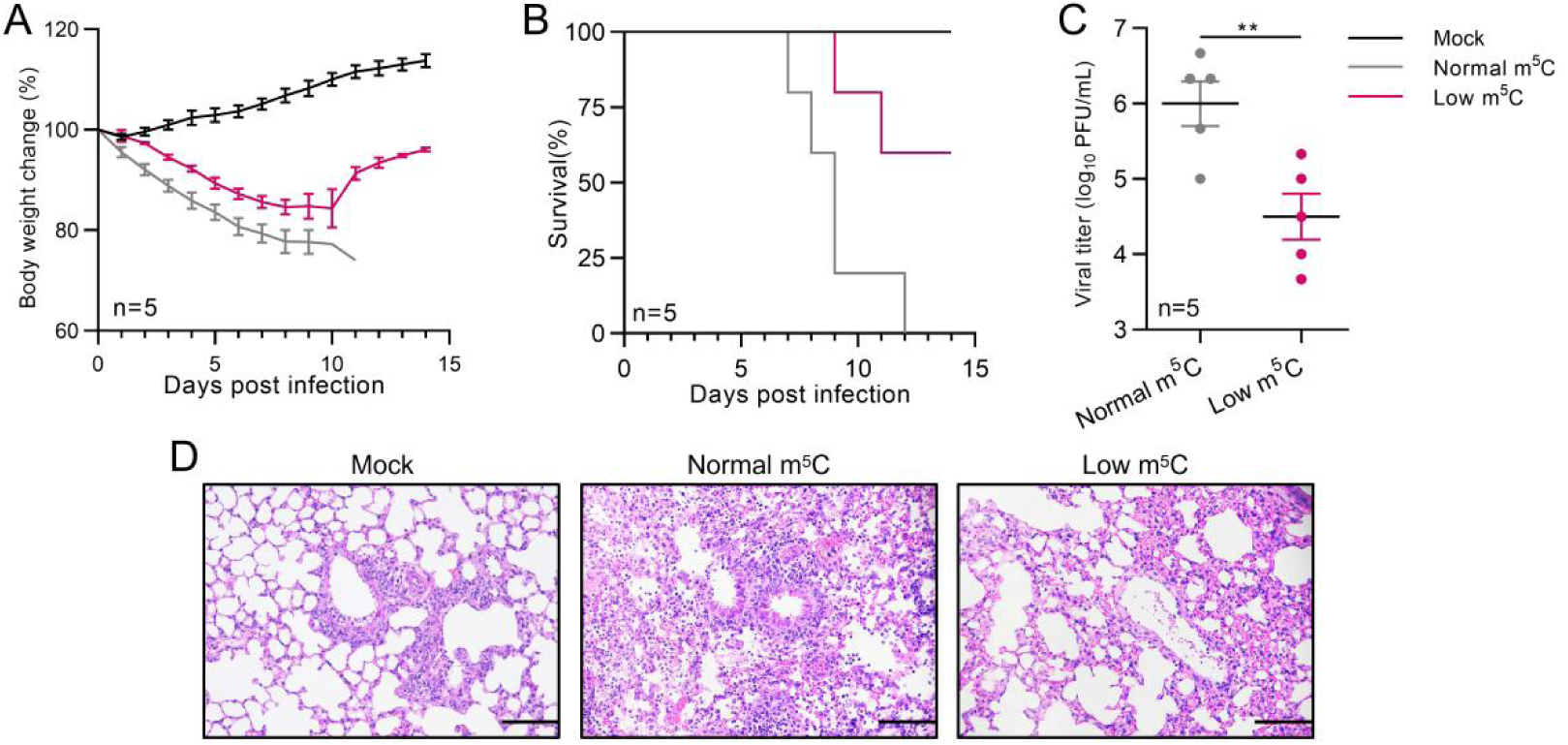
The low-m^5^C IAV has reduced pathogenicity in mice. (A) The body wight change curves of the IAV-infected mice. (B) The survival curves of the IAV-infected mice. (C) Viral titers measured in the lungs of infected mice at 5 dpi. Data are presented as mean ± SEM (n = 5). **, *P*<0.01 by two tailed *t*-test versus the WT. (D) Hematoxylin and eosin staining of lung sections of mice infected with the normal-m^5^C IAV and low-m^5^C IAV at 5 dpi. Each C57BL/6 mouse was infected with the indicated virus at a dose of 1 × 10^5^ PFU.

### Mutation of m^5^C modification sites on PB2 minus-strand RNA interferes with the selective packaging of IAV vRNPs

Multiple studies have demonstrated that the PB2 segment, encoding a subunit of viral polymerase, plays a critical role in the efficient and selective packaging of IAV vRNPs (14, 63–65). In this study, both the m^5^C MeRIP-seq and RNA-BisSeq results identified multiple m^5^C modification sites on the PB2 vRNA (Fig. 3), especially in the UTRs and nearby regions which are required for the genome packaging. To investigate the role of the m^5^C modification in the PB2 packaging regions in virus replication, we introduced synonymous mutations into the 5’and 3’ends of the coding region for *PB2* gene to replace the potential methylated C sites (position 144, 2178, 2187, 2190, 2199 and 2220 in the positive strand direction) in the PB2 vRNA, generating mutants WSN-vPB2_144_, WSN-vPB2_2178-2220_ and WSN-vPB2_144&2178-2220_ (*SI Appendix*, Fig. S5*A*). The reduction of methylation modification on the mutants were confirmed by MeRIP-qPCR assays, and WSN-vPB2_144&2178-2220_ containing all the six synonymous mutation sites had the lowest m^5^C level among the mutants (Fig. 6*A*). Subsequently, the replication ability of the mutant viruses was determined. In WT A549 cells, all the three mutants yielded lower titers of progeny viruses than the WSN-WT virus (Fig. 6*B*). While in *NSUN2*-KO cells, no significant differences in viral titers were observed between the WT and mutant viruses (Fig. 6*C*). Plaque assay also showed that the three PB2-vRNA mutants produced from A549 cells formed smaller plaques than the parental virus (Fig. 6*D*). However, by Western blot, mutation of m^5^C sites on the PB2 vRNA did not cause the reduction of viral protein production (*SI Appendix*, Fig. S5*B*). These results demonstrated that loss some of the m^5^C sites on PB2 vRNA did not impair the viral protein production but reduced the production of infectious virus particles, implying it may interfere with the assembly of infectious virus particles.

**Figure 6.**
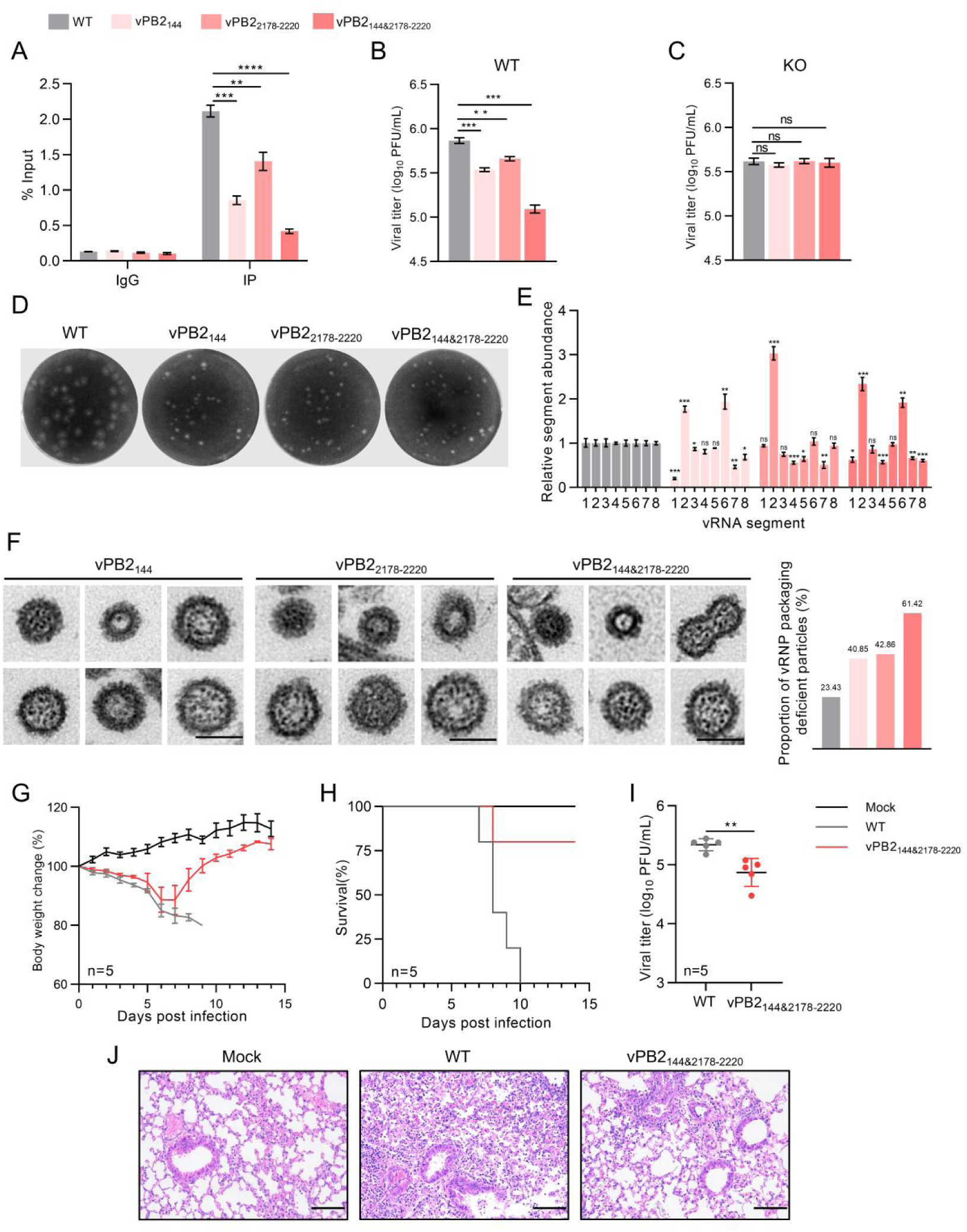
The m^5^C modification of PB2 minus-strand RNA regulates vRNA packaging. (A) Assessment of m^5^C level on WT and mutant PB2 vRNAs by MeRIP-qPCR. IP was performed using anti-m^5^C antibody, and IgG was used as a control antibody. (B) Titration of the indicated viruses produced in WT A549 cells. (C) Titration of the indicated viruses produced in *NSUN2*-KO A549 cells. (D) Plaque phenotype of WSN-WT and PB2-vRNA mutant viruses generated from A549 cells. (E) Relative abundance of the genome segments of PB2-vRNA mutants versus WSN-WT. 1-8: vRNA segments for PB2, PB1, PA, HA, NP, NA, M and NS. (F) Morphology of the PB2-vRNA mutants. Bars, 100nm. A particle with normal morphology is shown on the top left in each group, and others are abnormal particles. The percentage of abnormal virus particles is presented in a bar chart on the right. (G) The body wight change curves of the IAV-infected mice. (H) The survival curves of the IAV-infected mice. (I) Viral titers measured in the lungs of infected mice at 5 dpi. (J) Hematoxylin and eosin staining of lung sections of mice infected with the WSN-WT and WNS-vPB2_144&2178-2220_ at 5 dpi. Culture cells were infected with the indicated viruses at an MOI of 0.1. RNAs were extracted at 36 hpi and quantified by RT-qPCR. For plaque assay, viruses were harvested at 72 hpi. C57BL/6 mouse was infected with the indicated virus at a dose of 1 × 10^5^ PFU. Data are presented as mean ± SEM. *, *P*<0.05, **, *P*<0.01, ***, *P*<0.001 and ****, *P*<0.0001 by two-sided *t*-test versus the WT virus.

To examine the relative abundance of vRNA segments in the virus particles, vRNAs were extracted from WT and mutant IAV particles and quantified by RT-qPCR. The results showed that the relative abundance of PB1 fragment was significantly increased in the particles of all the three PB2-vRNA mutants, in accompany with the reduced packaging of several other vRNA fragments. The mutated PB2 fragment was significantly decreased in WSN-vPB2_144_ and WSN-vPB2_144&2178-2220_, while in these two mutant viruses, NA vRNA was also significantly increased besides the PB1 fragment (Fig. 6*E*). These data demonstrate that depletion of the m^5^C modification on PB2 results in an imbalance of the vRNA fragments packaged in IAV.

The morphology of the PB2-vRNA mutants was then examined by ns-TEM. Notably, the PB2-vRNA mutants not only form empty particles but also form some bigger-sized particles containing more than 8 vRNPs (Fig. 6*F*). Statistical data show that all the three mutant IAVs produce higher proportion of morphologically abnormal particles than the WT virus, with 40.85% (n = 213) for WSN-vPB2_144_, 42.86% (n = 231) for WSN-vPB2_2178-2220_, and 61.42% (n = 197) for WSN-vPB2_144&2178-2220_. The results of ns-TEM confirmed that mutation of m^5^C sites on PB2 vRNA severely interfered with the selective packaging of RNPs into IAV particles.

To investigate the pathogenicity of PB2-vRNA mutants, mice (n = 5) were infected with WSN-vPB2_144&2178-2220_ in parallel with WT virus at a does of 1×10^5^ PFU. The body wight of mice infected with WSN-vPB2_144&2178-2220_ decreased from the second day of infection and the survivals, which had reduced weight loss compared with the WT group, started to recovered at 7 dpi (Fig.6*G*). In two weeks, all the mice of WT group were died, in contrast, only one mouse infected with WSN-vPB2_144&2178-2220_ died (Fig.6*H*). At 5 dpi, the titers of WT virus in lungs were significantly higher than the WSN-vPB2_144&2178-2220_ group (Fig. 6*I*). Correspondingly, WSN-vPB2_144&2178-2220_ infection caused milder pathological changes in lungs than the WT virus (Fig. 6*J*). These data demonstrated that WSN-vPB2_144&2178-2220_ had reduced pathogenicity in mouse model.

## Discussion

As strict intracellular parasites, viruses hijack a large number of host factors to evade the host immune system and complete their replication cycles. Since the initial identification of 5-methylcytosine (m^5^C) modification in the viral RNA of Sindbis virus (32, 33) and adenovirus (34), this epigenetic mark added by host enzymes has also been detected in a number of other viruses including HIV-1 (27), MLV (39), SARS-CoV-2 (46), EV71 (44), HBV(35–37) and HCV (66). To date, RNA modifications including m^6^A, m^1^A, ac^4^C, m^7^G, inosine, and pseudouridine have been detected in IAV RNA (67–69). However, the presence and functional roles of m^5^C in IAV remained unexplored.

In this study, we mapped the m^5^C modification sites on the plus- and minus-strand RNAs of IAV by MeRIP-seq and RNA-Bis-seq, and found that most of the viral RNA fragments synthesized during IAV infection harbored m^5^C residues. Although the 5-methylcytosine modification of RNA is normally dynamic and reversible, and the addition and removal of m^5^C residues are coordinated by methyltransferases and demethylases (18, 70), some m^5^C sites in/close to the UTRs are consistently detected in the intracellular vRNA fragments and in the released virus particles. These regions highly overlap with the packaging signals of vRNAs that are required for genome incorporation and virion assembly.

Since NSUN2 has been known as the primary m^5^C writer for epitranscriptomic modification of various viral RNAs (27, 37, 39, 40, 44–46), we investigated whether NSUN2 participated in the m^5^C modification of IAV RNAs. We constructed NSUN2-C271A mutant able to crosslink with its target RNAs for m^5^C modification (27, 39) Detection of *NSUN2*-C271A in IAV particles supports that NSUN2 is involved in the modification of IAV RNAs. In *NSUN2* knockout cells, the m^5^C levels of IAV RNAs are significantly reduced, confirming that NSUN2 is an important m^5^C writer for IAV RNAs.

In retroviruses, NSUN2-mediated m^5^C has been shown to enhance viral gene expression and replication. Loss of NSUN2 impairs HIV-1 mRNA translation and splicing (27). NSUN2 also promotes MLV replication in an ALYREF reader protein-dependent manner (39, 40). Apart from retroviruses, positive-sense RNA viruses such as SARS-CoV-2, EV71 and HCV also utilize m^5^C modification to regulate viral RNA stability and replication (44–46). In our study, by changing the expression level of NSUN2 in A549 cells, we discovered that NSUN2 negatively regulates the production of IAV RNA and protein by affecting the transcriptional activity of vRNP rather than the stability of mRNA. Contrary to our results, m^5^C methylation has been reported to enhance retroviral and HBV mRNA translation (37). However, a recent study also report that overexpression of NSUN2 inhibites the secretion of HBV antigen, and knocking out of NSUN2 shows the opposite phenotype (35). It is still elusive how the NSUN2 or m^5^C residues regulate the transcription of vRNP.

In contrast to its downregulation role in viral gene transcription, m^5^C methylation on IAV vRNAs facilitated the generation of progeny virions. Interestingly, we also observed that the expression of NSUN2 was downregulated in early IAV infection, and then gradually upregulated after 36hpi. Previous studies have suggested that viral infections may downregulate NSUN2 as part of the host’s innate immune response (23, 24, 35), whereas viral proteins can also post-translationally upregulate methyltransferases (37, 45, 71, 72). It seems that NSUN2 expression is well tuned during IAV infection, being downregulated to promote the viral RNA and protein syntheses first, and then upregulated to facilitate the progeny virion assembly. The specific mechanism on how the NSUN2 expression is regulated needs to be further explored.

Selective packaging of the eight vRNAs into IAV particles is mediated by vRNA-vRNA and vRNA-NP protein interactions. Segment-specific packaging signals located in the 5’ and 3’ terminal coding and noncoding regions are indispensable for this process (59, 73). Each vRNA is assembled into a rod-shaped ribonucleoprotein (vRNP) with NP and the polymerase complex (PB2, PB1, PA), and eight vRNPs are organized into a “1+7” arrangement. It has been reported that deficiency of m^6^A modification inhibi<u>ts</u> vRNP assembly by impairing the interaction between vRNA and vRNP proteins (68). Here, our study revealed that NSUN2-mediated m^5^C modification is also required for the proper assembly of the segmented genome. Compared with the WT virus, the low-m^5^C IAV produced in *NSUN2*-KO cells and the IAV with synonymous mutations ablating several m^5^C modification sites in PB2 vRNA have more deficient virus particles with disordered vRNA incorporation and abnormal morphology. It is worth noting that the proportion of the three longest vRNAs (PB2, PB1 and PA) is significantly increased in the low-m^5^C virus particles harvested from *NSUN2*-KO cells. Meanwhile, the mutation of m^5^C sites on PB2 caused the elevation of the relative level of PB1 vRNA released with the virus particles. Using electron tomography, it has been observed that it is always one of the long vRNAs (PB2, PB1, PA or HA) occupies the central RNP position in the “1+7” structure (7). The increased proportion of the long vRNAs in low-m^5^C IAVs suggests that the central RNP may act as the primary factor triggering the assembly and encapsidation of the eight RNPs, and depletion of m^5^C on vRNAs may interfere with the assembly of the“1+7” structure and thus result in the formation of deficient IAV particles with imbalanced genome fragments.

At the RNA molecular level, m^5^C is known to stabilize RNA structures by promoting base stacking and increasing the thermal stability of hydrogen bonding with guanine (74, 75). In the structure of vRNP, some vRNA regions free of NP, which form secondary and/or tertiary structures that protrude from the twisted rod-like body of the vRNPs, likely play important roles in vRNA-vRNA interaction and selective genome packaging (76, 77). The m^5^C MeRIP-seq and RNA-BisSeq results in this study show that some IAV packaging signal regions are highly enriched with m^5^C residues. We speculate that the m^5^C modification on IAV vRNAs may participate in the virus assembly in several possible ways: 1. It may affect the direct recognition and binding between eight vRNA fragments, therefore influence the spatial arrangement of the eight vRNPs. 2. The disruption of m^5^C modification may influence the structure and interaction between RNPs via affect the vRNA-NP interaction. 3. The disruption of m^5^C modification could also influence the recruitment of other cellular and/or viral factors involved in the selective packaging of IAV genomes.

Viral m^5^C modification has been implicated in the pathogenicity of viruses. Ablation of this modification — either by mutating m^5^C sites in the viral genome or knockdown/out of the host m^5^C writer—significantly attenuates the virulence of VSV, RSV (23), HBV (37), HCV (45) and EV71 (44) in mice models. Consistent with these findings, our study demonstrated that the low-m^5^C IAV and PB2-vRNA mutant exhibit reduced replication and pathogenicity in mice. Similarly, mutation of m^6^A sites in IAV RNA reduces viral pathogenicity in mice (67, 68). These results highlight epigenetic modifications of IAV RNA as potential therapeutic targets for antiviral intervention.

In conclusion, our study provides the first comprehensive mapping and functional characterization of NSUN2-dependent m^5^C modifications in IAV RNA. m^5^C modification sites are mapped on both the plus and minus RNA strands of IAV, and the m^5^C modification machinery plays a dual regulatory role in the IAV replication cycle. The activity of NSUN2 repressed the IAV gene expression. Meanwhile, loss of NSUN2 and m^5^C modification on vRNAs interfered IAV assembly, resulting in the production of many deficient IAV particles which had a lower replication ability and pathogenicity in mice. These findings demonstrate that NSUN2-mediated m^5^C modification is a key regulatory factor in IAV biology and a potential target for novel anti-influenza strategies.

## Materials and Methods

### Mice and Ethics

C57BL/6 mice (6-week-old) were purchased from the SPF Laboratory Animal Center in Hubei University of Medicine. All animal experiments in this study were approved by the Research Ethics Committee of Hubei University of Medicine (approval No. HBMU-2022-063). The animal experiments were conducted strictly in accordance with the recommendations provided by the Guide for the Care and Use of Laboratory Animals of the Ministry of Science and Technology of the People’s Republic of China.

### Cells

Human A549 cell line was obtained from Shanghai Institute of Biochemistry and Cell Biology, and maintained in Ham’s F-12K (BasalMedia). Canine MDCK cell line was purchased from Cellcook Biotech, and maintained in Dulbecco’s modified Eagle’s medium (DMEM, BasalMedia). The cells were cultured in the media supplemented with 10% fetal bovine serum (FBS, Hyclone), 100 U/ml penicillin and 100 μg/ml streptomycin (Sangon Biotech) at 37℃ in a 5% CO_2_ incubator.

### Generation of gene knockout A549 cells

CRISPR-Cas9 was used to knock out *NSUN2* gene in A549 cells. The primers for the expression of a single guide RNA (sgRNA) targeting the *NSUN2* exon 2 were annealed and inserted into pX459 plasmid. All primers used in this study are listed in *SI Appendix*, Table S1. A549 cells transfected with the plasmids were selected with 2μg/ml puromycin. Single cell clones were obtained by limited dilution method. The depletion of NSUN2 in A549 cell was confirmed by Western blotting. The gene deletion was confirmed by sequencing analysis of the *NSUN2* exon 2 PCR fragment.

### Rescue of wild-type and recombinant IAVs

The 8-plasmids reverse genetics system was constructed from pPol plasmid. Each cDNA fragment from A/WSN/33(WSN, H1N1) was inserted into the vector between the human Pol promoter and mouse Pol terminator as described previously (78, 79). To rescue the virus, MDCK and 293T cells (1:5) were seeded in six-well plates, and transfected with the 8 plasmids (0.3μg each) when the cell monolayer reached up to 80% confluency. The medium was replaced with fresh DMEM containing 1% BSA, 0.25μg/μl TPCK-treated Trypsin (Sigma) at 24 h post-transfection (hpt). After a further 48 h, the culture supernatant was harvested. The rescued virus was passaged in MDCK cells and virus stocks were stored at -80℃.

To deplete m^5^C modifications on PB2 vRNA without changing the protein sequence and control elements for the transcription and translation of *PB2* gene, the cytosines on the vRNA at the nucleotide positions 144, 2178, 2187, 2190, 2199 and 2220 (in the positive strand direction) were synonymously mutated based on the result of m^5^C MeRIP-seq and RNA-BisSeq (*SI Appendix*, Fig. S5). The IAV mutants WSN-vPB2_144_, WSN-vPB2_2178-2220_ and WSN-vPB2_144&2178-2220_ were rescued using the 8-plasmid system.

### Plaque assay

MDCK cells were grown in 12-well plates to 80-90% confluency. Serial dilutions of the virus stock to be examined were added onto the MDCK monolayers and incubated at 37℃ for 1h. The media were then replaced with the mixture for plaque assay containing DMEM, 1% BSA, 0.25μg/μl TPCK-treated Trypsin, antibiotics and 1% low melting agarose (Sangon Biotech). Infected MDCK cells were incubated at 37℃ for 72h, stained with 1ml of 0.1% Neutral red, rinsed briefly with water, and allowed to dry at room temperature. The visible plaques were counted for the viral titer calculation.

### Hemagglutination (HA) assays

25μl of 1% chicken red blood cell (RBC) was added to 25μl of two-fold dilutions of virus in phosphate buffered saline (PBS), and the mixture was incubated at room temperature for 30 min. The HA titer was calculated as the reciprocal value of the highest virus dilution that resulted in complete hemagglutination.

### IAV infection *in vitro* and *in vivo*

A549 cells grown to 80%-90% confluence were infected with IAV at an MOI of 0.1, and incubated to allow the virus absorption at 37℃ for 2h. The medium was then replaced with F-12K supplemented with BSA, antibiotics and TPCK-treated Trypsin. The cells were incubated in 5% CO_2_ at 37℃. For animal infection, 6-week-old C57BL/6 mice were anaesthetized with isoflurane, and intranasally infected with IAV, or with an equal volume of PBS for mock infection. Five days after infection, five mice in each group were euthanized and lung tissues were taken for virus titration and Hematoxylin and eosin (H&E) staining. The body weight and survival numbers in each group (n = 5) were monitored consecutively for 14 days.

### Construction of NSUN2 expression plasmids and site-directed mutagenesis

*NSUN2* gene (NM_017755.6) was obtained by PCR using A549 cDNA as the template, then cloned into pcDNA3.1-FLAG, generating the NSUN2 expression vector pcDNA3.1-NSUN2-FLAG. C271A and C321A mutations in NSUN2 were introduced into the plasmid using ClonExpress II One Step Cloning Kit (Vazyme). All the constructs were verified by DNA sequencing.

### NSUN2 overexpression and rescue expression

For NSUN2 overexpression and rescue expression, A549 cells and *NSUN2*-KO cells were seeded in 12-well plates and then transfected with 1 µg of plasmid DNA (pcDNA3.1-NSUN2-FLAG or empty vector pcDNA3.1-FLAG) using lipo8000 (Beyotime). The transfected cells were infected with WSN at 24 hpt and harvested at the scheduled time points. The overexpression and rescue expression of NSUN2 protein were detected by Western blot.

### Detection of NSUN2 and its mutants packaged in virus particles

The expression plasmids for the expression of wild-type NSUN2, C271A or C321A mutants were transfected into *NSUN2-*KO A549 cells using lipo8000. At 24 hpt, the cells were infected with WSN isolate at an MOI of 0.1. At 72 hpi, the cell culture supernatant was harvested. To purify the virus particles released from *NSUN2-*KO cells, the supernatant was fractionated by 20-60% sucrose gradient ultracentrifugation. Alternatively, the virus particles were purified with a more stringent HAd technique (*SI Appendix*, Fig. S2*G*). The supernatant was mixed with RBCs to reach the final cell concentration of 0.2%, incubated with gently shaking at 4℃ for 30 min. The blood cells were then pelleted at 1250g for 5 min at 4℃, washed twice with chilled PBS, resuspended in 37℃ PBS and incubated at 37℃ for 30 min with gently shaking. Finally, the cells were pelleted by centrigugation at 1250g for 5 min at 25℃, and the virus particles released in the supernatant were lysed in SDS Loading Buffer and examined by Western blot.

### Western blot

Protein samples were lysed in 1×SDS Loading Buffer, resolved by 10% SDS-PAGE and transferred onto polyvinylidene difluoride (PVDF) membrane. After blocking with 5% skimmed milk overnight at 4℃, the membrane was incubated with a primary antibody, followed by incubation with an HRP-conjugated secondary antibody. Western blot signals were visualized by chemiluminescence. ImageJ was used to quantify the intensity of protein bands.

### RT-qPCR

Total RNA was isolated from IAV infected or uninfected cells with TRIzol reagent (Sangon Biotech) and isopropanol precipitation following the manufacturer’s protocol. cDNA was synthesized using HiScript III 1st Strand cDNA Synthesis Kit (Vazyme). All RT-qPCR experiments were performed using ChamQ Universal SYBR qPCR Master Mix (Vazyme) following the manufacturer’s instructions. GAPDH mRNA was used to normalize the RNA samples in RT-qPCR experiments. Each reaction mixture contained 2 μL of 10-fold diluted RT product, 10 μL of SYBR qPCR mix, and 4 μM segment-specific primers, in a final volume of 20 μL. Cycling conditions were 30 s at 95°C, followed by 40 cycles of 95°C for 10 s, 60°C for 30 s. The relative expression level of each RNA was calculated with the 2^−ΔΔCt^ method. All samples were analyzed in triplicate.

### mRNA stability assay

A549 cells and *NSUN2*-KO cells were seeded in a 12 well plate at a density of 1×10^5^ cells/well. On the second day, the cells were infected with IAV at an MOI of 1. At 5 hpi, 10 μg/ml Actinomycin D was added to inhibit mRNA transcription. Cells were harvested at 0, 4, 8, 12 and 16 h post the Actinomycin D treatment, and total cell RNA was extracted with TRIzol reagent. The mRNA was reverse transcribed using oligo dT primer, measured by qPCR.

### Detection of the primary transcription levels

To compare the primary transcription levels of the incoming vRNPs, A549 cells and *NSUN2*-KO cells seeded in 12-well dishes were infected with WSN virus at an MOI of 1, and maintained in the medium containing 100 μg/ml of cycloheximide (CHX). At 5 hpi, the mRNA was quantified by RT-qPCR.

### Determination of the proportion of the eight genomic vRNAs

Cells were harvested at 6 hpi after IAV infection (MOI=0.1). Total RNA was extracted using TRIzol reagent. To detect the proportion of vRNAs released in the supernatant, RNA was extracted at 60 hpi, reverse transcribed using vRNA specific primers (vRNA-RT), and quantified by qPCR respectively with segment-specific primers. Relative level of each vRNA was normalized to the level of PB2 vRNA, and then the percentage relative to the level of WT vRNA was calculated. Results are presented as the average data replicated in triplicate (77, 80).

### Reverse Transcription Droplet Digital PCR (RT-ddPCR)

The vRNA released in cell culture supernatant was extracted with TRIzol LS Reagent (Thermo Scientific). One-step RT-droplet digital PCR (ddPCR) Advanced Kit (Bio-Rad) was used to perform RT-ddPCR following the manufacturer’s instructions, on a Qx200® ddPCR System (Bio-Rad®). Thermal cycling conditions: reverse transcription at 42℃ for 60min, enzyme activation at 95℃ for 10min, followed by 40 cycles at 95℃ for 30 s and 60℃ for 1 min, and then enzyme deactivation at 98℃ for 10 min. Each reaction was set up in triplicate, and the data were collected from three independent experiments.

### Electron microscopy

Wild-type or *NSUN2-*KO A549 cells grown in 10 cm dishes were infected with IAV or the mutant viruses at an MOI of 1. At 18 hpi, the cells were harvested and washed with pre-cooled 1×PBS. The cells were fixed with 2.5% glutaraldehyde for 2h at room temperature, with fresh 2.5% glutaraldehyde on ice for another 2 h, followed by fixation in 2% osmium tetroxide. Then the cells were sequentially dehydrated with 50%, 70%, 90%, 95%, and 100% ethanol, block-mounted and thinly sliced. Images were obtained by transmission electron microscopy (Hitachi, HT7800).

### m^5^C MeRIP-qPCR and MeRIP-seq

To examine IAV mRNAs, total RNA was extracted from infected A549 cells at 36hpi. To examine IAV vRNAs, the RNA was extracted from the cell culture supernatant which was harvested at 60 hpi and then concentrated by PEG6000. m^5^C MeRIP-qPCR was performed as previously described with some modifications (20). One-tenth of each RNA sample was kept as input control. 2 μg of anti-m^5^C antibody (Epigentek, 33D3) or IgG antibody (Sangon, D110503) was incubated with 25 μl protein G (Beyotime) in 50 μl IP buffer (10 mM Tris–HCl pH 7.5, 150 mM NaCl, 0.1% Igepal CA-630 (v/v)) for 1h at room temperature, followed by 3 rinses with 200 μl IP buffer. Then RNA was added and incubated with the antibody immobilized on beads for 1h at 4℃. The RNA-antibody-beads complexes were washed three times with 200 μl IP buffer and then incubated in 100 μl elution buffer (5 mM Tris–HCl pH 7.5, 1 mM EDTA, 0.05% SDS, and 80 μg proteinase K) for 1 h at 50°C, repeating the elution step for five times. RNA was precipitated from the eluate using ethanol and sodium acetate. The eluted RNA and input control RNA were reverse-transcribed with strand-specific reverse primers and quantified by qPCR.

To map the m^5^C modification profile of IAV RNA, total RNA was extracted from the virus-infected A549 cells at 36hpi. The MeRIP-seq analysis was performed by DIATRE Biotech. Briefly, m^5^C RNA immunoprecipitation was conducted using GenSeqTM m^5^C RNA IP Kit (GenSeq Inc). Both the input samples and the m^5^C IP samples were used for the construction of RNA-seq libraries using NEBNext^®^ Ultra II Directional RNA Library Prep Kit (New England Biolabs). The library quality was evaluated with BioAnalyzer 2100 system (Agilent Technologies). Library sequencing was performed on an illumina NovaSeq 6000 instrument with 150bp paired-end reads. For data analysis, adaptor sequences and low-quality reads in fastq files were removed using Cutadapt software with default settings (85). The clean reads of all libraries were aligned to the reference genome (IAV, A/WSN/33) by Hisat2 software (v2.0.4) to generate SAM files (86), which were then processed by Samtools to perform sorting and indexing, and then ultimately converted into BAM files (87). Subsequently, the bam files were separated into positive and negative strand alignments based on the orientation of the reads during library preparation. Finally, MACS2 was used to identity the possible m^5^C peaks on the plus and minus strands of the virus (88).

### RNA-BisSeq

The RNA released in cell culture supernatant was isolated using TRIzol reagent. The RNA-BiS libraries were constructed by E-GENE Co. Ltd as previously described (81, 82). Briefly, the extracted RNA was subjected to DNase I (NEB) treatment at 37℃ for 30 min to remove residual DNA. Bisulfite conversion of the isolated RNA samples was conducted using EZ RNA Methylation Kit (Zymo Research) at 70 ℃ for 5min, followed by incubation at 54℃ for 45 min. The converted RNA sample was purified using Zymo-Spin IC Column according to the manufacturer’s instructions. A non-methylated lambda DNA sequence was added as a control during library construction, and the bisulfite conversion efficiency of this sequence was higher than 99.5% in all the samples to make sure that the non-methylated C sites were effectively converted. Reverse transcription was carried out using ACT random hexamers and Superscript II Reverse Transcriptase (Invitrogen), and the synthesized cDNA was then used for library construction. After analyzed by Agilent 2100 Bioanalyzer (Agilent Technologies) and quantified by real-time PCR, RNA-BS libraries were finally sequenced using Illumina Novaseq 6000 platform (Illumina) with paired end 150-bp read length.

Low quality bases and Adapter sequences were trimmed using Trimmomatic (v0.38) (83). The cleaned reads were aligned with WSN reference genome using BS-RNA (v1.0) (84). The vRNA sites with methylation rate (m^5^C/C%) ≥1% in two independent experiments are considered as potential modification sites.

### Statistical analysis

Data were shown as the mean ± SEM and the significance of differences was evaluated by two tailed t-tests between two groups. The *p* values are marked as follows: ns (not significant) means *P* > 0.05, \**P* < 0.05, \*\**P* < 0.01, \*\*\**P* < 0.001, \*\*\*\**P* < 0.0001.

---

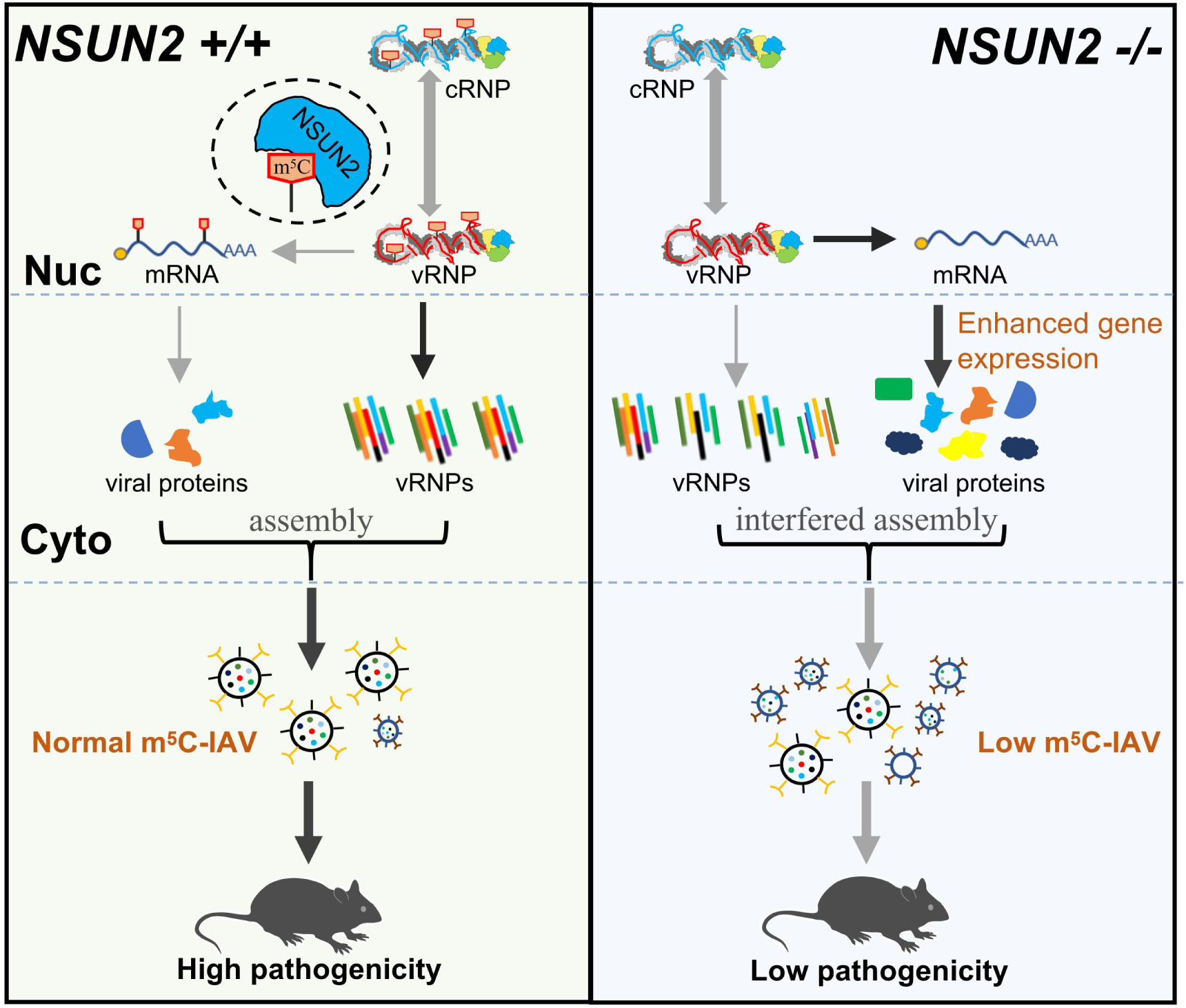
Schematic model for the roles of NSUN2-dependent m^5^C modification in IAV gene expression and genomic RNA packaging. m^5^C is highly enriched in plus and minus IAV RNA strands and the methyltransferase NSUN2 is an important m^5^C writer. Although NSUN2 negatively regulates viral RNA synthesis and protein translation, its methytransferase activity is required for the efficient packaging of IAV segmented genome. Loss of m^5^C modification on vRNAs interferes IAV assembly, resulting in the production of many deficient IAV particles which have a lower replication ability and reduced pathogenicity in mice.

