## Supplemental Figure and Table for "NSUN2-dependent 5-methylcytosine Modification Regulates Influenza A virus Gene Expression and Genomic Packaging"

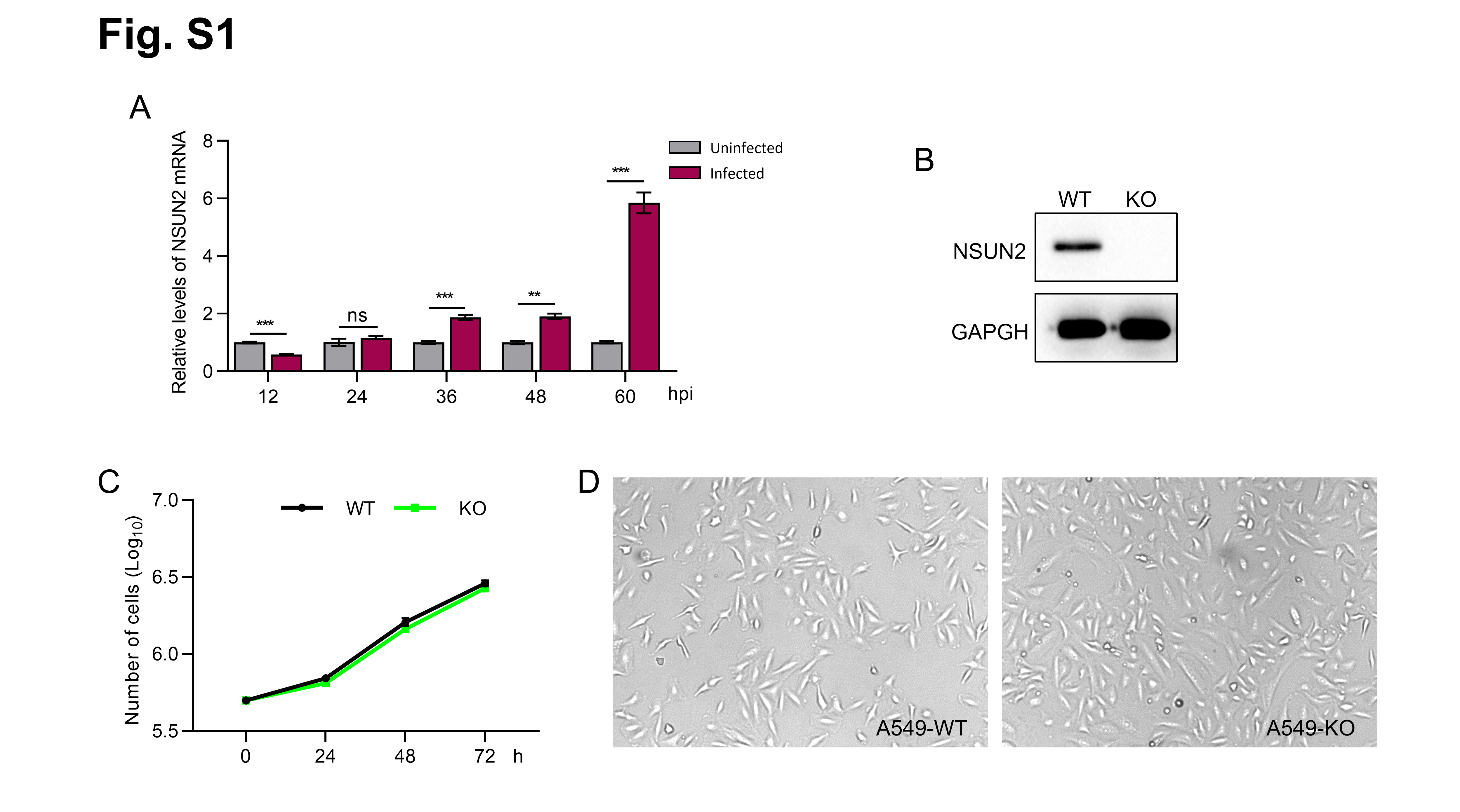

**Fig. S1. Effects of IAV infection on NSUN2 expression and characterization of the *NSUN2*-KO A549 cell line.**

(*A*)The NSUN2 mRNA levels measured by RT-qPCR during IAV infection of A549 cells. Data are presented as mean ± SEM (n = 3). **, P<0.01, ***, P<0.001 and ns, not significant by two-sided t-test versus the uninfected sample.

(*B*) Confirmation of NSUN2 depletion by Western blot.

(*C*) The growth curves of wild-type and *NSUN2*-KO A549 cells, n = 3.

(*D*) The cell morphology of wild-type and *NSUN2*-KO A549 cells.

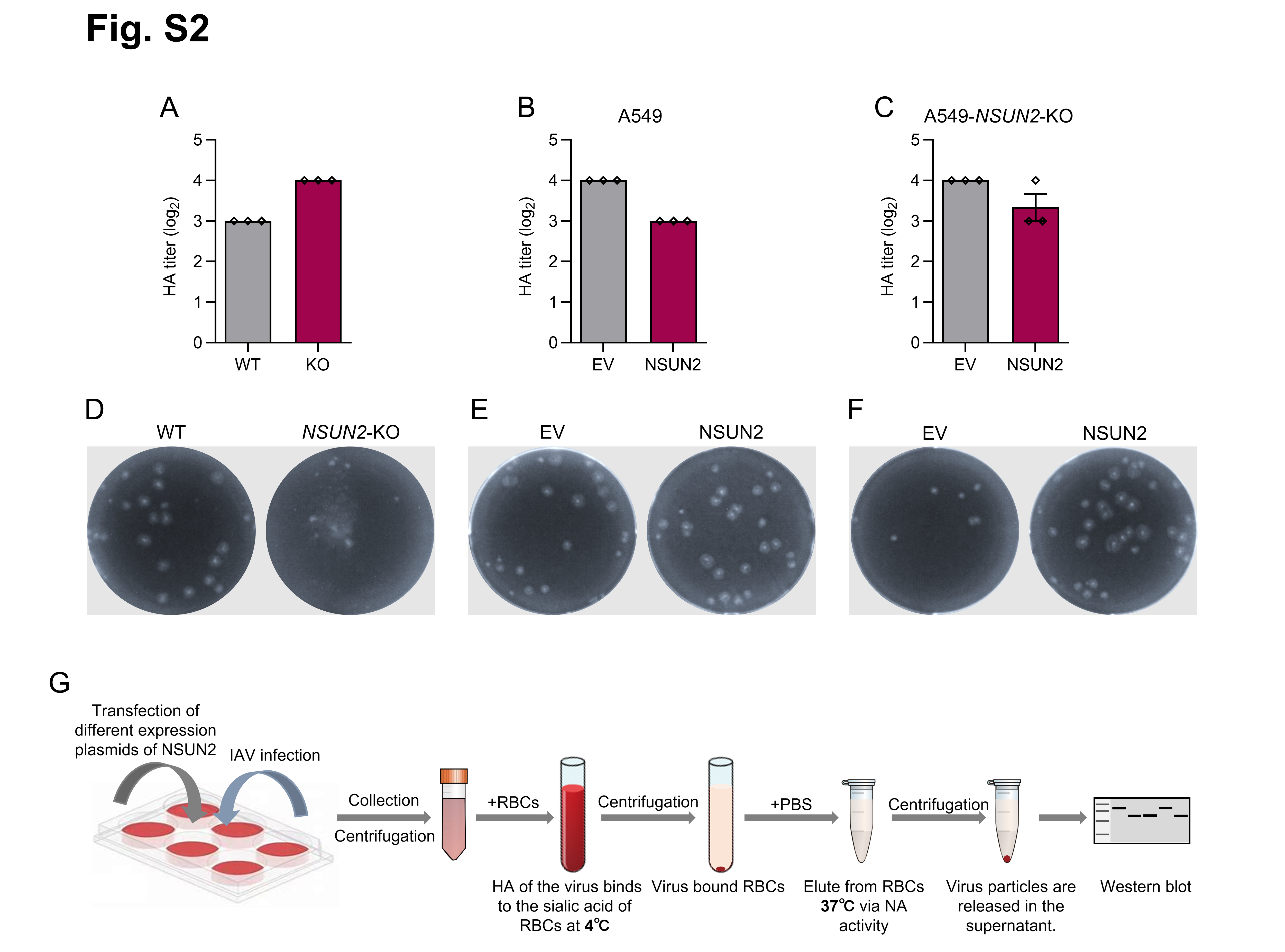

**Fig. S2. NSUN2 positively regulates the production of IAV progeny virus.**

(*A*) Depletion of NSUN2 resulted in an increase of HA titer.

(*B*) Overexpression of NSUN2 caused the drop of HA titer in IAV-infected A549 cell medium.

(*C*) Rescue expression of NSUN2 in *NSUN2*-KO A549 cells suppressed the increase of HA titer.

(*D*) to *(F*) Plaque assays of IAV stocks harvested in (*A*) to (*C*)*.*

(*G*) Schematic diagram of the protocol for purification of IAV particles by haemadsorption/elution (HAd). RBCs: red blood cells.

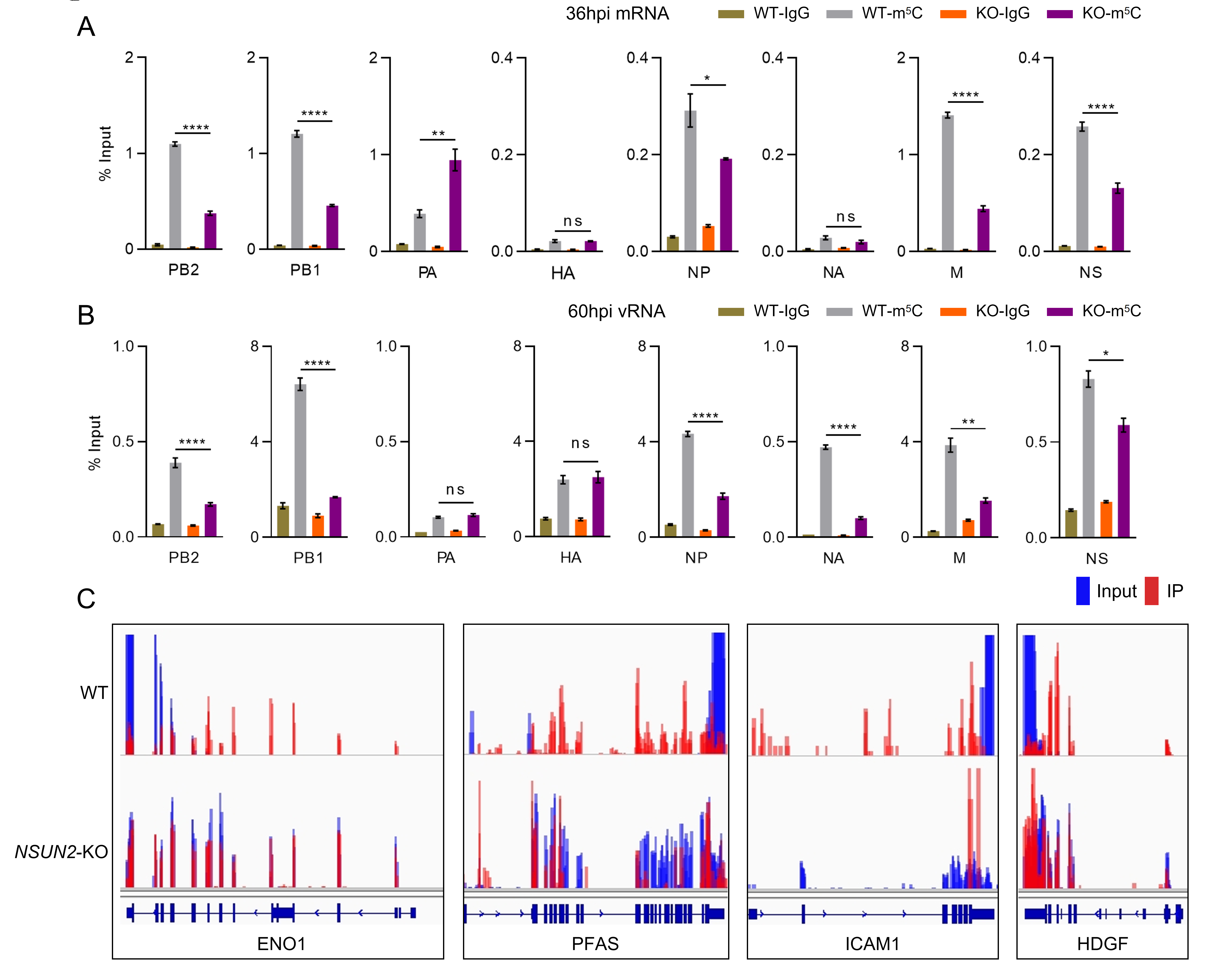

**Fig. S3. MeRIP-qPCR detection of NSUN2-mediated m^5^C modification of IAV mRNA and genomic RNAs and analysis of control host transcripts in m^5^C MeRIP-seq.**

(*A*) and (*B*) Assessment of the m^5^C modification on IAV mRNAs *(A*) and vRNAs (*B*) by MeRIP-qPCR. A549 cells and *NSUN2*-KO A549 cells were infected with IAV-WSN at an MOI of 0.1. mRNA was extracted from infected cells at 36 hpi, and vRNA was extracted from the cell culture supernatant at 60 hpi. Methylated RNAs were retained by m^5^C antibody immobilized on magnetic beads, and then quantified by RT-qPCR. Mouse IgG was used as a control antibody in the assay. Data are presented as mean ± SEM (n = 3). *, *P*<0.05, **, *P*<0.01, ***, *P*<0.001, ****, *P*<0.0001 and ns, not significant by two-sided *t*-test versus the WT group.

(*C*) The m^5^C modifications of the indicated four transcripts which have been reported to be catalyzed by NSUN2 (1-4) are analyzed as the positive controls for MeRIP-seq in this study.

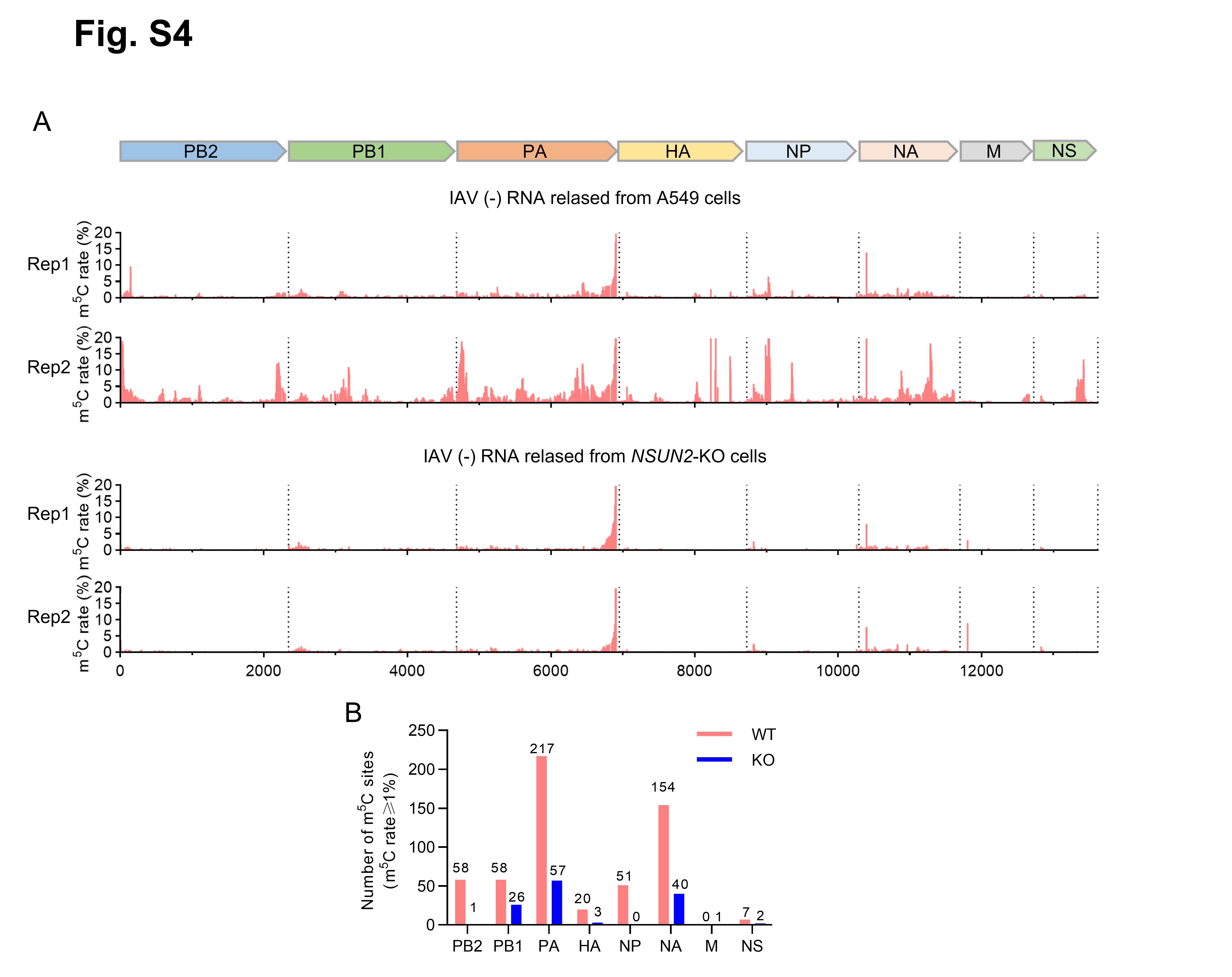

**Fig*.* S4.** m^5^C sites on IAV vRNAs identified by RNA-BisSeq.

WT and *NSUN2*-KO A549 cells were infected with IAV-WSN at an MOI of 0.1. At 60 hpi, the virus particles released in the cell culture supernatant were used for RNA extraction and subjected to RNA-BisSeq. The concatenated map of the IAV-WSN genome is shown on the top for the alignment of detected methylation sites. Data for two independent experiments are shown. A non-methylated lambda DNA sequence was added as a control during library construction, and the bisulfite conversion efficiency of this sequence was higher than 99.5% in all the samples to make sure that the non-methylated C sites were effectively converted.

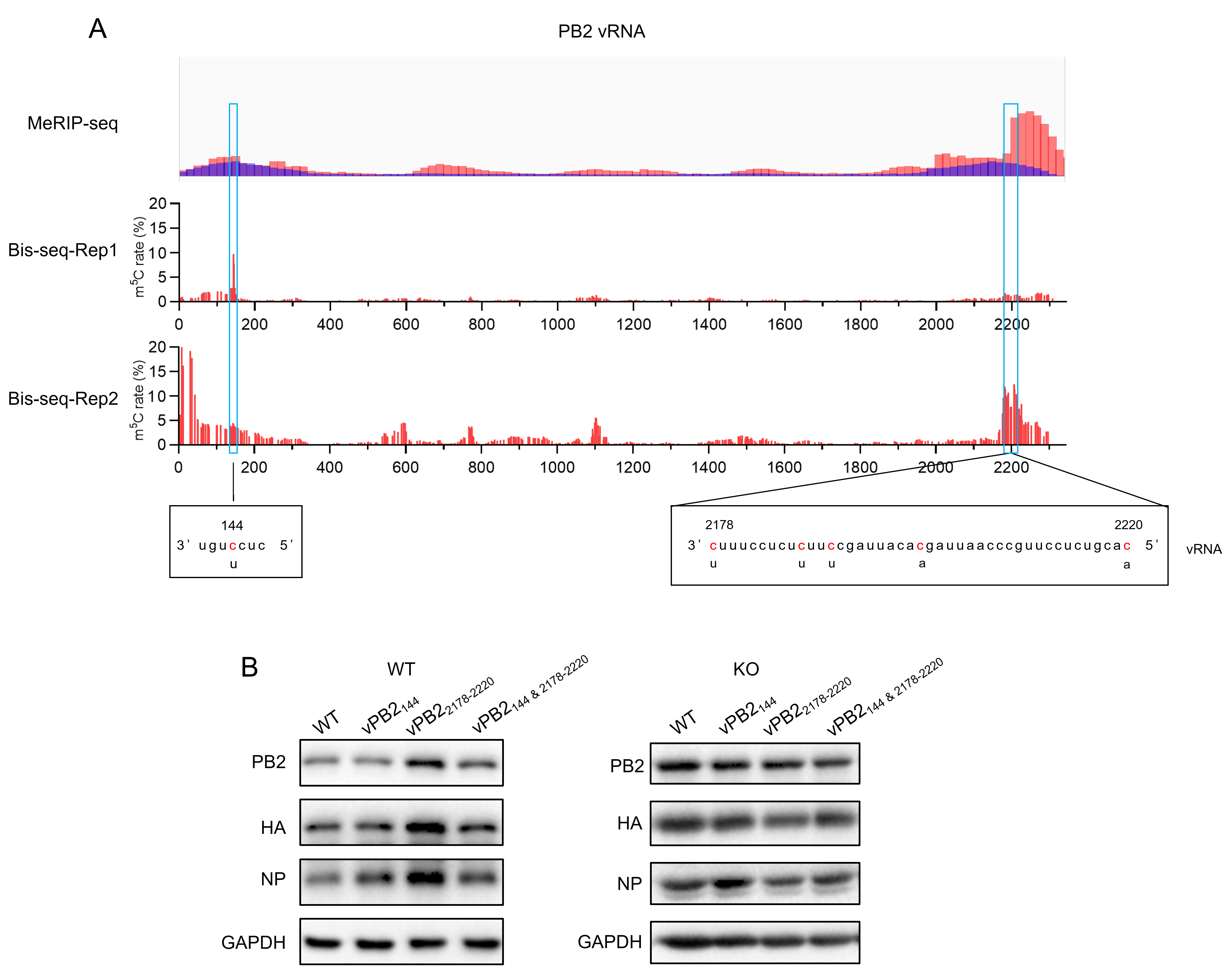

**Fig. S5. Synonymous mutation of m^5^C sites on PB2 vRNA.**

(***A***) The mutation sites on PB2-vRNA mutants. The top IGV track displays the MeRIP-seq, in which the IP results are shown in red and the Input reads are shown in blue. The RNA-BisSeq results show the ratio of unconverted C sites on PB2 vRNA. The blue boxes highlight the modification regions detected both in MeRIP-seq and RNA-BisSeq, and can be synonymously mutated. The mutation sites and sequences at positions 144, 2178, 2187, 2190, 2199 and 2220 (position numbered in the positive strand direction) are shown in the black boxes.

(*B*) Viral protein levels detected by Western blot. WT and *NSUN2*-KO A549 cells were infected with the indicated viruses at an MOI of 0.1, and the whole cell lysates were harvested at 24 hpi for Western blot analysis.

Table S1. Key resources table

| REGENT or RECOURSE | SOURCE | IDENTIFIER |
| --- | --- | --- |
| Antibodies | | |
| Anti-NSUN2 | Proteintech | Cat# 20854-1-AP |
| Anti-GAPDH | TransGen | Cat# HC301 |
| Anti-PB2 | GeneTex | GTX125926 |
| Anti-NP | GenScript | Custom-produced |
| Anti-WSN serum | This Study | N/A |
| Anti-m^5^C | Epigentek | Cat# 33D3 |
| Mouse IgG | Sangon | Cat# D110503 |
| Anti-Mouse IgG (HRP) | CWBIO | Cat# CW0102S |
| Anti-Rabbit IgG (HRP) | CWBIO | Cat# CW0103S |
| Bacterial and Virus Strains | | |
| IAV-WSN | This Study | N/A |
| WSN-mNP_247-723_ | This Study | N/A |
| WSN-vPB2_144_ | This Study | N/A |
| WSN-vPB2_2178-2220_ | This Study | N/A |
| WSN-vPB2_144 & 2178-2220_ | This Study | N/A |
| Top10 | This Study | N/A |
| Chemicals and Animal blood product | | |
| Lipo8000 | Beyotime | Cat# C0533 |
| Puromycin | Beyotime | Cat# ST551 |
| BSA | Solarbio | Cat# H1130 |
| FBS | VivaCell | Cat# C04001 |
| TPCK-treated Trypsin | Sigma | Cat# T1426 |
| Agarose, Low melting | Sangon | Cat# A600015 |
| RNase Inhibitor | Beyotime | Cat# R0102 |
| Ribonucleoside Vanadyl Complexes, RVC | Beyotime | Cat# R0107 |
| Proteinase K | Beyotime | Cat# ST532 |
| 1% Chicken Erythrocyte  Cycloheximide  Actinomycin D | Sbjbio  AbMole  AbMole | Cat# SBJ-RBC-C001  Cat# M4879  Cat# M4881 |
| Critical Commercial Assays | | |
| Total RNA Extractor | Sangon | Cat# B511311 |
| HiFiScript gDNA Removal cDNA Synthesis Kit | CWBIO | Cat# CW2582 |
| 2 × Rapid Taq Master Mix | Vazyme | Cat# P222 |
| 2 × Phanta Max Master Mix | Vazyme | Cat# P515 |
| ChamQ Universal SYBR qPCR Master Mix | Vazyme | Cat# Q711 |
| BeyoMag™ Protein G Magnetic Beads | Beyotime | Cat# P2105 |
| Experimental Models: Cell lines | | |
| A549 | This Study | N/A |
| MDCK | cellcook | Cat# CC9504 |
| A549-*NSUN2-*KO | This Study | N/A |
| Experimental Models: Organism/strain | | |
| Mouse | the SPF Laboratory Animal Center | Strain: C57BL/6 |
| Oligonucleotides | | |
| Oligo name | Sequence | |
| sgNSUN2-fwd | caccgtgttctccttgacgatctcg | |
| sgNSUN2-rev | aaaccgagatcgtcaaggagaacac | |
| vRNA-RT | AGCGAAAGCAGG | |
| cRNA-RT | AGTAGAAACAAGG | |
| mRNA-RT(oligo dT) | TTTTTTTTTTTTTTT | |
| GAPDH-RT | GAAGATGGTGATGGGATTTC | |
| NSUN2-qPCR-fwd | gctggcacaggagggaatata | |
| NSUN2-qPCR-rev | TGCCAGGTCCTTTGCTTGAc | |
| GAPDH-qPCR-fwd | ctacactgagcaccaggtgg | |
| GAPDH-qPCR-rev | catgaggtccaccaccctg | |
| PB2-qPCR-fwd | ATGTGAGGGGATCAGGAATG | |
| PB2-qPCR-rev | GCCTTCATCTGGGTCTTCAG | |
| PB1-qPCR-fwd | ACCCACTGAACCCATTTGTC | |
| PB1-qPCR-rev | GATCCAGGAGTGTGTTGTTG | |
| PA-qPCR-fwd | AGAGCCTATGTGGATGGATTCG | |
| PA-qPCR-rev | TTGGACCGCTGAGAACAGG | |
| HA-qPCR-fwd | GCGAACAACTCAACCGACAC | |
| HA-qPCR-rev | TGGAAGCAGTGAGTCGCATT | |
| NP-qPCR-fwd | CAGGATGTGCTCACTGATGC | |
| NP-qPCR-rev | TTCTCCGTCCATTCTCACCC | |
| NA-qPCR-fwd | GAATCGGTTGCTTGGTCAGC | |
| NA-qPCR-rev | GGGCCATCGGTCATTATGGT | |
| M-qPCR-fwd | ATGAGAACCGTTGGGACTC | |
| M-qPCR-rev | TGCAATGACGAGAGGATCAC | |
| NS-qPCR-fwd | GCATCGCGCTACCTAACTGA | |
| NS-qPCR-rev | CCATGATCGCCTGGTCCATT | |
| Recombinant DNA | | |
| pcDNA3.1-Flag | This Study | N/A |
| pcDNA3.1-NSUN2-Flag | This Study | N/A |
| pcDNA3.1-*NSUN2-*C321A-Flag | This Study | N/A |
| pcDNA3.1-*NSUN2-*C271A-Flag | This Study | N/A |
| pX459 | MiaoLing | Cat# P52786 |
| Deposited Data | | |
| m^5^C MeRIP-seq  (H1N1-infected A549 cells) | HRA003296 | https://ngdc.cncb.ac.cn/gsa-human/s/0HTMk5a3 |
| m^5^C MeRIP-seq  (H1N1-infected *NSUN2*-KO A549 cells) | HRA006236 | https://ngdc.cncb.ac.cn/gsa-human/s/a3h87T0t |
| RNA-BisSeq  (IAV released from A549 cells) |  |  |
| RNA-BisSeq  (IAV released from *NSUN2*-KO A549 cells) |  |  |
| Software and Algorithms | | |
| Cutadapt | Martin (5) | https://cutadapt.readthedocs.io/en/stable/ |
| Hisat2 | Kim et al.(6) | http://daehwankimlab.github.io/hisat2/ |
| Samtools | Li et al.(7) | http://samtools.sourceforge.net |
| MACS2 | Zhang et al. (8) | https://pypi.org/project/MACS2/ |
| IGV | Robinson et al. (9) | http://software.broadinstitute.org/  software/igv/ |
| Trimmomatic  BS-RNA | Bolger et al. (10) | https://github.com/usadellab/Trimmomatic |
|  | Liang et al. (11) | https://ngdc.cncb.ac.cn/bsrna/ |
| GraphPad prism | Graphpad software | https://www.graphpad.com/features |
| ImageJ  RNAfold | N/A  N/A | <https://imagej.nih.gov/ij/>  http://rna.tbi.univie.ac.at/cgi-bin/RNAWebSuite/RNAfold.cgi |
